# Androgen stimulation rapidly reorganizes temporal 3D genome and epigenome states to trigger AR-mediated transcription in prostate cancer

**DOI:** 10.1101/2024.12.17.628805

**Authors:** Elyssa M. Campbell, Geraldine Laven-Law, Alicia Jones, Dayna Challis, Timothy J Peters, Grady Smith, Yolanda Colino-Sanguino, David Moulder, Qian Du, Katherine A. Giles, Daniel W. Thomson, Clare Stirzaker, Theresa A. Hickey, Fatima Valdes Mora, Amanda Khoury, Susan J. Clark, Joanna Achinger-Kawecka

## Abstract

Androgen receptor (AR)-mediated transcription drives prostate cancer progression and remains a critical therapeutic target. AR activation by androgens triggers nuclear translocation, DNA binding at AR regulatory elements (AREs) and transcription initiation. This process involves co-factors such as FOXA1, epigenetic modifications, and 3D chromatin interactions. However, the temporal coordination of these events during AR-mediated transcription remains poorly understood.

Using a time-course androgen stimulation model in prostate cancer cells, we integrated temporal multi-omics data within the MOFA+ framework to dissect AR-mediated transcriptional dynamics. Our analysis revealed that rapid AR binding at gene promoters is crucial for initiating nascent transcription of AR target genes. We identified H3K27 acetylation at flanking AREs, pre-marked by constitutive FOXA1 binding, occurs prior to AR recruitment. Additionally, androgen stimulation induced a temporal early shift in nascent transcription from MYC to AR target genes, associated with transient disruption of 3D chromatin interactions. Finally, we showed that CRISPR-mediated inhibition of AR-associated chromatin contacts suppresses distal linked target genes, demonstrating that dynamic genome architecture is functionally required for AR-driven transcription.

Together these findings uncover the dynamic reorganization of 3D chromatin structure and transcription factor binding post-androgen stimulation, resulting in distinct temporal epigenomic states. This work provides new insights into the mechanisms governing AR-mediated transcriptional activation in prostate cancer and highlights the interplay between MYC and AR in regulating transcription in hormone-driven cancers.

## INTRODUCTION

Androgen receptor (AR)-mediated transcription is the primary driver of prostate cancer progression and remains the key therapeutic target for treatment. While AR responsiveness is a key factor in discerning successful treatment strategies, its dynamic nature and evolution to complex genomic states independently of AR, often results in the development of resistance to AR-targeted drugs^1^. AR in its inactive form resides in the cytoplasm, but when activated by androgens, such as testosterone or 5α-dihydrotestosterone (DHT), AR translocates into the nucleus where it binds to DNA at AR binding sites (AR regulatory elements; AREs) to initiate transcription^2^. This process involves a multifaceted interplay of coregulatory proteins and pioneer transcription factors, including FOXA1, that work in concert with AR binding at regulatory DNA elements to induce the transcription of AR target genes^3^. The activation of AREs is associated with epigenetic and chromatin features, including transcription factor binding and post-translational histone modifications^4,5^. The majority of AREs are located distal from gene promoters^6^ where they act as enhancers^7^, and are brought into close physical proximity by three-dimensional (3D) chromatin interactions^8^. Enhancers are proposed to induce transcriptional changes through long-range 3D chromatin interactions and recruitment of co-regulatory proteins and transcriptional machinery^9,10^. However, the direct contribution of chromatin contacts to gene regulation is less clear^11,12^, as studies that removed chromatin loops genome-wide report conflicting results^13,14^. Moreover, the potential multifaceted and dynamic interplay of epigenetic modifications, 3D chromatin interactions and transcription factor binding involved in the temporal activation of AR-mediated transcription has not been addressed.

To investigate how induction of AR binding at DNA regulatory elements, epigenome modifications, co-regulator recruitment and 3D chromatin interactions together might contribute to AR-mediated transcription, we generated a time-course experimental model of androgen stimulation in prostate cancer cells and integrated the temporal multi-omics datasets into a MOFA+ framework. Notably, our time-resolved multi-omics approach reveals an underappreciated role of rapid AR binding at gene promoters which contribute to nascent transcription of AR-mediated genes. We demonstrate that androgen stimulation, within just 5 minutes, induces rapid AR binding at AREs, that are pre-marked by acetylation of H3K27 histones and by constitutive FOXA1 binding.. Furthermore, we reveal a very early change in nascent transcription from MYC to AR target genes that occurs immediately post androgen stimulation, and propose this is mediated by transient androgen-induced 3D chromatin interactions at these regulatory elements. Together, our results clarify the temporal dynamics of epigenomic reprogramming during AR-mediated transcriptional activation in prostate cancer cells.

## MATERIAL AND METHODS

### Prostate cancer cell culture

LNCaP (RRID: CVCL_0395) and LAPC-4 (RRID: CVCL_4744) prostate cancer cells were maintained in RPMI-1640 medium (Gibco #11875093; LNCaP) or Iscove’s Modified Dulbecco’s Medium (Gibco #12440061; LAPC-4) supplemented with 10% (v/v) Fetal Bovine Serum (FBS, Sigma # F9423), as described previously^15^. All cell lines were authenticated by STR profiling (CellBank Australia, Westmead, NSW, Australia) and cultured for <6 months after authentication. Mycoplasma contamination testing was routinely performed in the cell lines while in culture (MycoAlert™ Mycoplasma Detection Kit, Lonza). Cells were seeded at 80% confluence and cultured for 3 days in phenol red-free RPMI-1640 medium (Gibco #11835055; LNCaP) or Iscove’s Modified Dulbecco’s Medium (Gibco #21056023; LAPC-4) supplemented with 5% charcoal-stripped FBS with daily washing to thoroughly deplete cells of residual androgens. After 3 days growth, cells were, stimulated with 10 nM 5α-dihydrotestosterone (DHT; Sigma #A8380; CAS #521-18-6) for 5 mins, 10 mins, 20 mins, 30 mins, 2 hrs, 4 hrs and 16 hrs. For the vehicle samples, the cells were treated with ethanol (Sigma #E7023) to a final of 0.001%.

### Chromosome conformation capture with *in situ* Hi-C

Hi-C in LNCaP cells was performed using the Arima-HiC kit, according to the manufacturer’s protocols (Arima HiC Cat. #A510008, Arima Capture-HiC+ Kit, cat#A510008, cat#A303010 and cat#A311025). Briefly, cells were cross-linked with 2% formaldehyde to obtain 1-5 μg of DNA per Hi-C reaction. The Arima kit uses two restriction enzymes: ^GATC (DpnII), G ^ANTC (HinfI), which after ligation of DNA ends generates 4 possible ligation junctions in the chimeric reads: GATC-GATC, GANT-GATC, GANT-ANTC, GATC-ANTC. Hi-C libraries were prepared using the KAPA Hyper Library Prep Kit (Roche, cat#KK8502) or Arima Library Prep kit (cat# A303010) with a modified protocol provided by Arima with 10 PCR cycles for library amplification as required.

### Promoter Capture Hi-C (PCHi-C)

Promoter Capture Hi-C (PCHi-C) in LNCaP cells was performed as previously described ^16^ using the Arima HiC+ Capture kit with the Promoter Capture Panel (#Cat A302010). Promoter Capture was carried out using Arima Hi-C libraries derived from two replicates of LNCaP cells with the SureSelect target enrichment system and RNA bait library according to manufacturer’s instructions (Agilent Technologies SureSelect XT HS2 DNA Reagent Kit cat#5190-9685) or with Arima Capture-HiC+ Kit (cat#A510008, cat#A303010 and cat# A311025), using 12 post-capture PCR cycles as required. PCHi-C libraries were sequenced on the Illumina NovaSeq S4 platform at the Garvan Sequencing Platform.

### Chromatin Immunoprecipitation with Sequencing (ChIP-seq)

ChIP-seq^17^ in LNCaP and LAPC-4 cells was performed in three independent biological replicates (sequential passages of cells), as described previously^18^. Chromatin was prepared using the BioRuptor Plus (Diagenode) and immunoprecipitated with either 5 µg of AR) antibody (Abcam #Ab108341) or 2 µg of H3K27ac antibody (Abcam #Ab4729) coupled to Protein A Dynabeads (Invitrogen #1002D). Each H3K27ac ChIP (ChIP-Rx) was calibrated via spike-in with ∼10% *Drosophila melanogaster* chromatin prepared from Schneider’s Drosophila Line 2 cells (Thermo Fisher #R69007; CVCL_Z232). ChIP DNA was either submitted for library preparation at the South Australian Genomics Centre (SAGC) using the Qiagen Ultra-Low DNA Library Prep Kit (cat#180497), or prepared in-house using the NEBNext Ultra II DNA Library Preparation Kit (New England Biolabs #E7645). Libraries were sequenced at the SAGC to a depth of >20 M reads per sample, either on the Illumina NextSeq 550 System (1x 75 bp High-Output flow cell, cat#20024906) or on the DNBSEQ-T7 (2 x 150 bp, MGI #940-000268) after DNA nanoball conversion.

### Cleavage Under Targets and Release Using Nuclease (CUT&RUN)

The CUTANA™ ChIC/CUT&RUN Kit (EpiCypher, 14-1048) was used to perform the CUT&RUN assay as described previously^19^ with minor modifications. CUT&RUN experiments included antibodies targeting AR, c-MYC, FOXA1 (Abcam, cat#Ab108341, cat# Ab32072 and cat#Ab23738), H3K27ac (Thermo Fisher Scientific, cat#MA5-23516) H3K27me3 and CTCF (EpiCypher, cat#13-0055 and cat#13-2014). Briefly, 150,000 cross-linked LNCaP cells were input per reaction and three independent biological replicates were generated per treatment time-point. Cells were lightly cross-linked for 1 minute with 0.1% formaldehyde according to manufacturer’s recommendations. Cross-linked cells were then quenched with 2.5M glycine for 5 minutes, scraped and washed with PBS and 1x Protease Inhibitors, followed by the standard protocol. Prior to the final clean-up of CUT&RUN DNA, reverse crosslinking was done by incubating overnight at 55-degrees with 10% SDS and 20μg/μl Proteinase K. Library preparation was performed using the CUTANA™ CUT&RUN Library Prep Kit (EpiCypher, 14-1001 and 14-1002) as per the manufacturers protocol. Libraries were pooled and sequenced on the Illumina NovaSeq S4 6000 system (2 x 150bp reads).

### mRNA-seq

Total RNA was extracted from three biological replicates of LNCaP cells using TRIzol reagent (Invitrogen, cat#15596026) and purified using ethanol precipitation. Sequencing libraries were prepared with TruSeq Stranded poly-A RNA Library Prep kit (Illumina) by Ramaciotti Centre for Genomics (UNSW Sydney, Australia). Libraries were sequenced on Illumina NovaSeq S4 in paired-end mode.

### TT-Seq

TT-Seq protocol performed based on a previously published method^20^. Briefly, 1mM 4sU was added to cell culture media for the last 5 minutes of each DHT treatment time-point. RNA isolation was done using 3mL of TRlzol Reagent (Invitrogen, cat#15596026) and 5PRIME Phase Lock tubes (Heavy Gel, Quanta, Cat#2302830) using the required cell numbers to yield ∼400µg of RNA (∼20x10^6^ LNCaP cells/sample). RNA was sheared on a Covaris M220 Focused-Ultrasonicator for 4 seconds with the following parameters: peak power 50W, 20% duty factor, 200 cycles per burst at 7°C. After sheering, RNA was diluted to 130ng/µL in a buffer, maintaining the following ratio 38uL RNA: 1uL 1M HEPES: 1uL 0.5M disodium EDTA. The 4sU-labelled fraction of RNA was biotinylated with MTSEA-biotin-XX at 50ug/ml in dimethylformamide (DMF) for 30 minutes with continuous rotation (tube rotator, 18rpm 45-degrees fixed rotation). uMACS columns (1x per 100ug per sample) were used to separate 4sU-labelled RNA (Miltenyi Biotec, µMACS™ Streptavidin Kit cat#130-133-282 and µ columns cat#130-042-701). The Ramaciotti Centre for Genomics (UNSW Sydney, Australia) performed sample QC, rRNA depletion, library preparation (Illumina Stranded Total RNA-seq with Ribo Zero Plus, cat# 20040525), and sequencing on the NovaSeq X (2x100bp).

### Total RNA-Seq

Following separation of 4sU-labelled RNA for TT-seq, the unlabelled non-nascent fraction of RNA was purified by ethanol precipitation, and Total RNA-Seq libraries were prepared with by Ramaciotti Centre for Genomics (UNSW Sydney, Australia) in the same way as the 4sU-labelled TT-Seq samples.

### CRISPRi

dCas9-KRAB lentivirus was produced from a second-generation lentiviral vector expressing a KRAB-dCas9 fusion protein (pHR-SFFV-KRAB-dCas9-P2A-mCherry, Addgene #60954, RRID:Addgene_60954) in HEK293T cells (RRID #CVCL_0063) transfected with packaging plasmids as per standard protocols. dCas9-KRAB lentiviral supernatant was delivered to LNCaP cells, and successfully transduced cells were selected on mCherry expression using the FACSAria III Cell Sorter (BD BioSciences). Guide RNAs (gRNAs) targeting AR binding sites, the *KLK2* promoter, and a non-targeting control (NTC) (**Supplementary Table 1**) was designed using CRISPOR and cloned lentiviral CRISPR guide vector pJR85 (Addgene #140095; RRID:Addgene_140095) using the BstXI/BlpI cut sites. LNCaP dCas9-KRAB cells were further transduced with guide RNA lentivirus (representing 1 – 3 different guide RNAs) at MOI 1 and maintained in culture medium supplemented with 2 µg/ml puromycin (Invivogen #ANT-PR-1). Both pHR-SFFV-KRAB-dCas9-P2A-mCherry and pJR85 CRISPRi plasmids were a kind gift from Jonathan Weissman.

### RT-PCR

Prostate cancer cells were collected into TriReagent (Sigma #T9424). All RT-PCR experiments were performed in at least three independent biological replicates (sequential passages of cells). Total RNA was extracted from cell lysates using the Direct-Zol RNA kit (Zymo #R2052), and reverse transcribed with the SuperScript III First-Strand Synthesis Kit (Invitrogen #18080400) and random hexamers. Quantitative RT-PCR was performed on the CFX384 Real-Time PCR Detection System (BIO-RAD) with PowerTrack SYBR Green Master Mix (Applied Biosystems #A46113) paired with primers outlined in **Supplementary Table 1**. Gene expression was calculated using the 2^-ΔΔCt^ method and normalised *GAPDH* expression. Where indicated, CRISPRi data was further normalised to gene expression from the CRISPRi Non-Targeting Control.

### Immunoblotting

Prostate cancer cells were scraped into radioimmunoprecipitation assay buffer and debris removed via syringing and centrifugation. Protein concentration was quantified by BCA protein assay (Thermo Fisher #23227) and immunoblotting was performed using standard techniques. The ChemiDoc (BIO-RAD) Imaging System was used to visualise (1) total protein and (2) PSA expression after overnight incubation in primary antibody (Cell Signalling Technology, #5365T, 1:500) followed by secondary antibody incubation (Cell Signalling Technology. #7074S) and development with ECL Select (Cytiva #2235). PSA expression was quantified by band densitometry using ImageLab (BIO-RAD) software for two experimental replicates, representing sequential passages of cells.

### Promoter Capture Hi-C analyses

The sequenced reads were processed using HiCUP (v.0.9.2)^21^ and mapped to the hg38/GRCh38 genome digested with the –arima flag and with the minimum di-tag length set to 20. Unique, valid mapped reads from HiCUP were converted into .chinput files using bam2chicago.sh utility, and obtained .chinput files were further filtered and processed with CHiCAGO (v.1.14.0)^22^. CHiCAGO design files were created based on the Arima-HiC design of the bait fragments: MaxLBrowndist, 75,000; binsize, 3,000; minFragLen, 25; maxFragLen, 1,200. Significant interactions were called with CHiCAGO using a score cut-off of five. All bait-to-bait interactions were discarded. Chicdiff package (v.0.6)^23^ was used to compare PCHi-C data between the different time-points. Interactions were initially pre-filtered based on between-replicate reproducibility (replicate SD < Q3) and mean CHiCAGO score in the top 5000 per sample group. Chicdiff was then used to identify differential chromatin interactions from the top 5000 interactions per sample group in pairwise comparisons with the EtOH-treated control. For visualization of the significant interactions in WashU Epigenome Browser^24^, replicates were merged with CHiCAGO.

### ChIP-seq data analyses

Sequenced ChIP-Seq data was aligned to the hg38/GRCh38 reference genome with bowtie2^25^ (version 2.2.9), using default parameters. Mapped reads were sorted and indexed using Samtools^26^ (version 1.11). Peak calling was performed with MACS2 ^27^ (version 2.2.7.1) using the callpeak function with default parameters to call narrowPeaks. Deeptools^28^ (version 3.5.0) bamCoverage function was used to call bigwig signal, with reads in the ChIP-Seq blacklisted regions from ENCODE ^29^ excluded. Differential binding analysis of AR ChIP-Seq data was done with the R package, DiffBind^30^, and comparisons were performed pairwise between each DHT treatment time-point and the EtOH treated control. Differential AR binding was determined for each comparison by an absolute logFC > 0.6 and FDR < 0.05. AR ChIP-seq data for LNCaP cells treated in an independent DHT experiment was downloaded from GEO (GSE251898) to compare to the datasets generated in the study (Supplementary Note). All H3K27ac ChIP data were calibrated via additional alignment to the *Drosophila* dm6 reference genome. Briefly, the total dm6-aligned read counts were calculated with samtools and was used as a scaling factor for both downstream bigwig signal visualization. H3K27ac DiffBind analysis was performed with spike-in normalisation.

### CUT&RUN data analyses

Paired-end fastq files were down-sampled to to 10 million uniquely mapped read pairs per sample using seqtk (https://github.com/lh3/seqtk) and aligned to the hg38/GRCh38 reference genome using the Bowtie v.2 algorithm using the nf-core “cutandrun” pipeline^19,31^. Merged bigwig tracks for visualization were created from merged bam files from all replicates using the bamCoverage function with scaling factor normalization, and heatmaps and average profiles were plotted with deepTools^28^.

### mRNA-seq data analyses

RNA-seq raw reads were quality controlled and sequence adaptors were trimmed using Trim Galore (v.0.11.2). Reads were mapped with STAR (v.2.7.7a)^32^ to the hg38/GRCh38 human genome build with GENCODE v.33 used as a reference transcriptome in RSEM^33^ (parameter settings: –quantMode TranscriptomeSAM–outFilterMatchNmin 101 –outFilterMultimapNmax 20). TMM normalization was applied to normalize for RNA composition and differential expression was performed with edgeR v.3.18.1^34^ using the GLMQLF with FDR < 0.05 and absolute logFC > 1. RNA-seq tracks were generated using bedtools v.2.22 genomeCoverageBed to create normalized.bedGraph files and bedGraphToBigWig (USCS utils) to create .bigwig files.

### TT-seq data analyses

TT-seq and Total RNA raw reads were quality controlled and sequence adaptors were trimmed using Trim Galore (v.0.11.2). Reads were mapped with STAR (v.2.7.7a)^32^ to the hg38/GRCh38 human genome build with GENCODE v.33 used as a reference transcriptome in RSEM^33^ (parameter settings: –quantMode TranscriptomeSAM–outFilterMatchNmin 101 –outFilterMultimapNmax 20).

EdgeR was used to calculate normalization factors with TMM normalization per sample group, including replicate TT-Seq and Total RNA-Seq libraries with the same treatment and DHT time-point. Differential nascent expression analysis was performed per sample group with edgeR v.3.18.1^34^ using the GLMQLF with FDR < 0.05 and absolute logFC > 1. Signal tracks were generated using bedtools v.2.22 genomeCoverageBed to create normalized.bedGraph files and bedGraphToBigWig (USCS utils) to create .bigwig files. ChromHMM^35^ was used to define the nascent and steady RNA states by training a 4-state model from TT-Seq and Total RNA-Seq signals per DHT time-point with the EtOH treatment as a control. Size factor normalised coverage in 200bp bins using fragment midpoints was computed as follows. TT:Total RNA-seq nascent expression signal tracks were generated using deeptools bamCoverage function with parameters to centre piled-up reads on the midpoint of bam fragments. A 200bp bin size and normalization factors from edgeR as scale factors were used and reads per genome coverage (RPGC) normalisation was performed per strand. Finally, nascent expression was quantified as the log2 ratio of these TT-Seq to Total RNA-Seq signals.

### Enrichment analyses

The HOMER motif discovery suite (v.4.10)^36^ was used for motif analysis, using default background. Motifs were ranked by log *P* values from hypergeometric enrichment calculations (or binomial) to determine motif enrichment. Observed over expected enrichment was analyses were performed using GAT^37^ and ChIPenrich. Unique, non-redundant ChIP-seq binding sites from the ReMap22 database^38^ were used to identify potential co-binding events. Peak annotation analyses were performed using ChIPPeakAnno and ChIPSeeker.

### Gene ontology analyses

Gene ontology enrichment analysis and pathway enrichment were done using GSEA (v.4.1.0)^39^ and MSigDB 7.2^40^. All significant biological processes and pathways had an adjusted P value of <0.001.

### MOFA temporal variance modelling

The statistical framework from MOFA^41^ was used to model the temporal variance in differential data sets with DHT treatment (16-hour time course). MEFISTO^42^ is a probabilistic factor analyses model implemented as part of the MOFA framework and is available as Bioconductor package MOFA2 (version 1.3.3). To prepare for model training, three differential data sets (AR ChIP-seq, mRNA-seq and PCHi-C) were prepared. Training of the model was carried out using the following options: training three factors with DHT treatment time as the covariate of interest, Automatic Relevance Determination prior on the factors = False, Automatic Relevance Determination prior on the weights = True, Spike-and-slab prior on the factors = False, Spike-and-slab prior on the weights = True and gaussian likelihoods per view. By default, MEFISTO uses a squared exponential kernel with Euclidean distances to define a covariance function. The trained MOFA model represented data variability in terms of three latent factors, which were further explored and visualized.

### Statistical analyses

The Mann–Whitney–Wilcoxon test was used for two-group non-parametric comparisons. Unless otherwise stated, statistical tests were two-sided. A permutation test (n = 1,000 permutations) was used to calculate empirical *P* values in observed/expected analyses. The Benjamini–Hochberg method was used to control for multiple testing using an FDR procedure. Different mean values per group were compared using a two-tailed unpaired Student’s t-test and plotted using ggpubr package (https://rpkgs.datanovia.com/ggpubr/index.html).

### Code availability

Python script language (v.2.7.8 and v.3.9.1) and R (v.4.0.3) were used to develop the bioinformatics methods and algorithms in this work. All code for PCHi-C, MOFA+ and TT-seq analyses is available within the GitHub repository (https://github.com/elyssa-parbery/AR_3DEpigenome_PrCa).

## RESULTS

### Establishment of a temporal integrated molecular model of androgen stimulation

To model the temporal dynamics of AR-mediated transcription in prostate cancer following androgen deprivation, we performed a time-course of androgen DHT or vehicle (EtOH control) treatment on LNCaP hormone-sensitive prostate cancer cells with three replicates and collected cells at four different time-points (30-minutes, 2-hours, 4-hours, and 16-hours) of treatment (**Fig. 1A**). First, to validate AR binding was induced by DHT treatment in a temporal manner, we performed AR ChIP-Seq and analysed differential binding with DHT time-points in comparison to the EtOH treated control. Clustering of biological replicates and separation of treatment conditions demonstrated robust and dynamic gain in AR binding associated with DHT stimulation (**Fig. 1B**). Using DiffBind^44^, we identified 10,950 AR binding sites that were gained after 30-minutes of DHT treatment, compared to EtOH, 15,279 AR binding sites after 2-hours, 17,375 AR binding sites after 4-hours and 20,074 AR binding sites after 16-hours of DHT treatment (**Fig. 1C** & **Supp. Fig. 1A**). The majority (54.5%) of androgen-induced AR binding occurred at the first time-point of 30-minutes and these sites were maintained until the last tested time-point (16-hours) (**Fig. 1D**).

**Figure 1:**
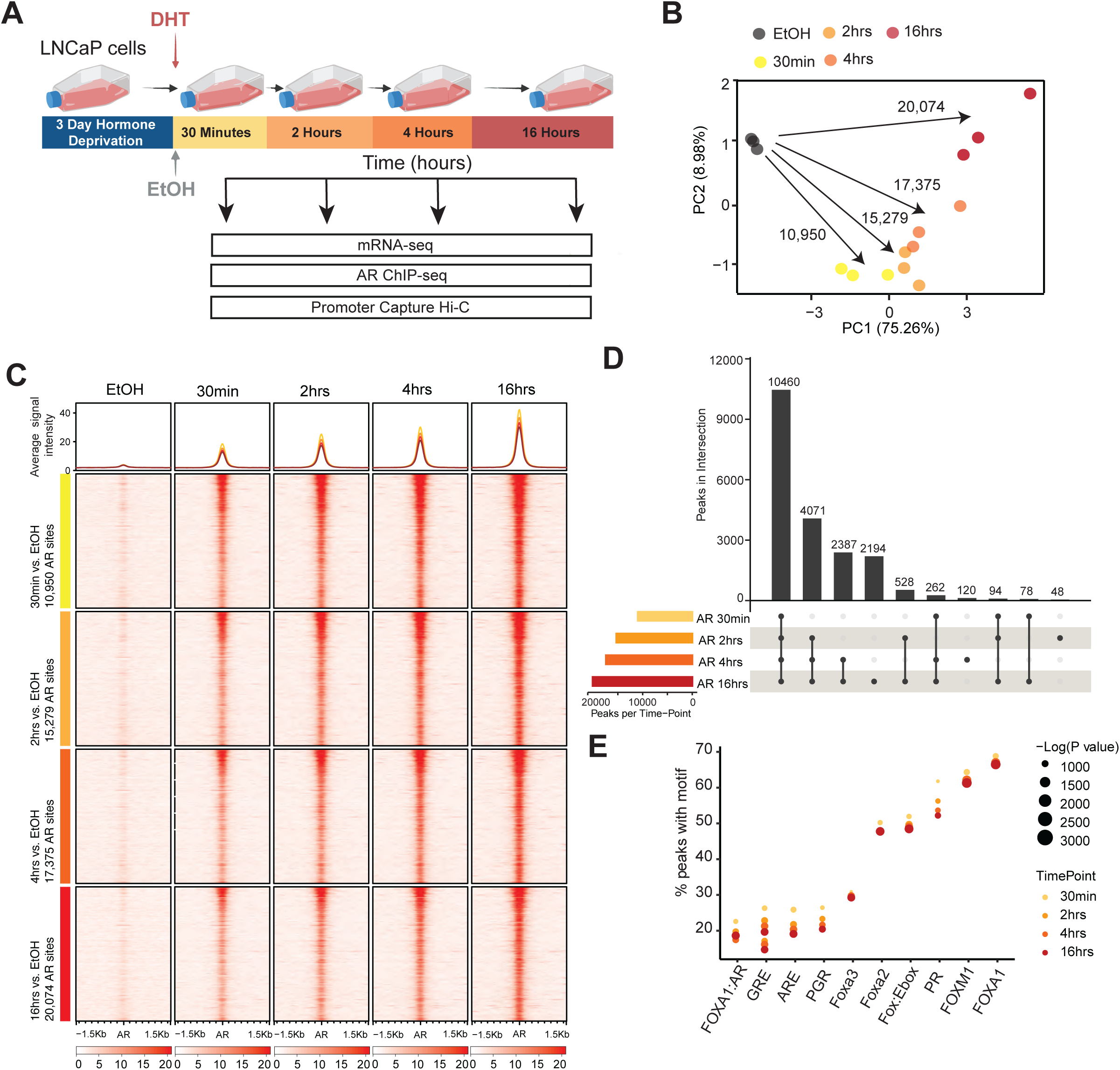
Temporal model of androgen-stimulation in LNCaP cells. **(A)** Schematic overview of the experimental design. LNCaP cells were treated with 10nM DHT and samples were collected at five time-points for the androgen-stimulation time-course (30mins, 2hrs, 4hrs, 16hrs and EtOH (control)) for mRNA-seq, AR ChIP-seq and Promoter Capture Hi-C (PCHi-C). **(B)** Principal component analyses showing the relationship among the AR binding ChIP-seq profiles upon DHT stimulation in the androgen-stimulation time-course. Data plotted for normalised read counts. The number of differentially bound AR peaks between each DHT time-point and EtOH is shown. **(C)** Heatmap and average profiles of AR ChIP-Seq (n=3 merged replicates) signal intensity at AR binding sites for differentially bound AR peaks for each time-point of DHT stimulation compared to EtOH control. **(D)** Overlap of differentially bound AR peaks at each time-point of DHT stimulation compared to EtOH control presented in an upset plot. **(E)** Transcription factor motifs enriched at differentially bound AR peaks (FDR < 0.05 and logFC > 0.6) identified by two-way analyses between each DHT treatment time-point and EtOH control. Percent of AR peaks with motif was compared to matched random regions across the genome.

We next overlapped the differential AR binding sites at each time-point with the different regulatory regions across the genome using LNCaP ChromHMM states^10^ and performed transcription factor (TF) motif enrichment analysis. The highest percentage of AR binding sites gained after 30-minutes of DHT treatment were located at active enhancers (41.4%), followed by active promoters (23.1%) and weak enhancers (17.1%) (**Supp. Fig. 1B**). This distribution and significant fold change enrichment for different ChromHMM states (**Supp. Fig. 1C**) was maintained for AR binding sites gained at 30-minutes, 2-hours, 4-hours, and 16-hours, consistent with the large overlap between gained AR binding across treatment time-points. In addition, using HOMER^36^ we identified that the gained AR peaks across the DHT treatment time course were also significantly enriched for FOX TF motifs (FOXM1, FOXA1, FOXA2, FOXA3, Fox:Ebox and FOXA1:AR), GRE Nuclear receptor (NR) and ARE (NR) TF motifs, and progesterone receptor (PR) (**Fig. 1E**).

Next, to interrogate the temporal order of AR transcription activation, we characterized gene expression changes by performing matched RNA-seq across the DHT treatment time-course. Principal component analysis (PCA) revealed a trajectory along which hormone-deprived LNCaP cells (EtOH, control) acquired androgen-responsive gene expression across the time course (**Fig. 2A**). We performed differential gene expression (DGE) analyses at each time point to discover if there is a temporal androgen-responsive gene expression pattern. In total, we found expression of 1,921 unique genes were significantly altered (false discovery rate (FDR) <0.01) between any time point of DHT treatment and EtOH (**Fig. 2A**); 122 (74 UP and 48 DOWN) differentially expressed genes at 30min of androgen treatment, 793 (622 UP and 171 DOWN) at 2-hours, 659 (432 UP and 227 DOWN) at 4-hours and 806 (527 UP and 279 DOWN) at 16-hours of androgen treatment (**Supp. Fig. 2A**, **Supplementary Table 2**). Most gene expression changes occurred at the 2-hour time-point (793/1921; 41.3%) and 16-hour time-point (806/1921; 41.9%), however a large proportion (1899/1921; 98.8%) of differential gene expression was not maintained through the DHT treatment time-course (**Supp. Fig. 2B**). We found that differentially expressed genes frequently overlapped gained AR binding sites at their promoters (**Supp. Fig. 2C)** and AR binding proximal to gene promoters (< 1kb) was associated with increased expression across the time-course (**Supp. Fig. 2D**).

**Figure 2:**
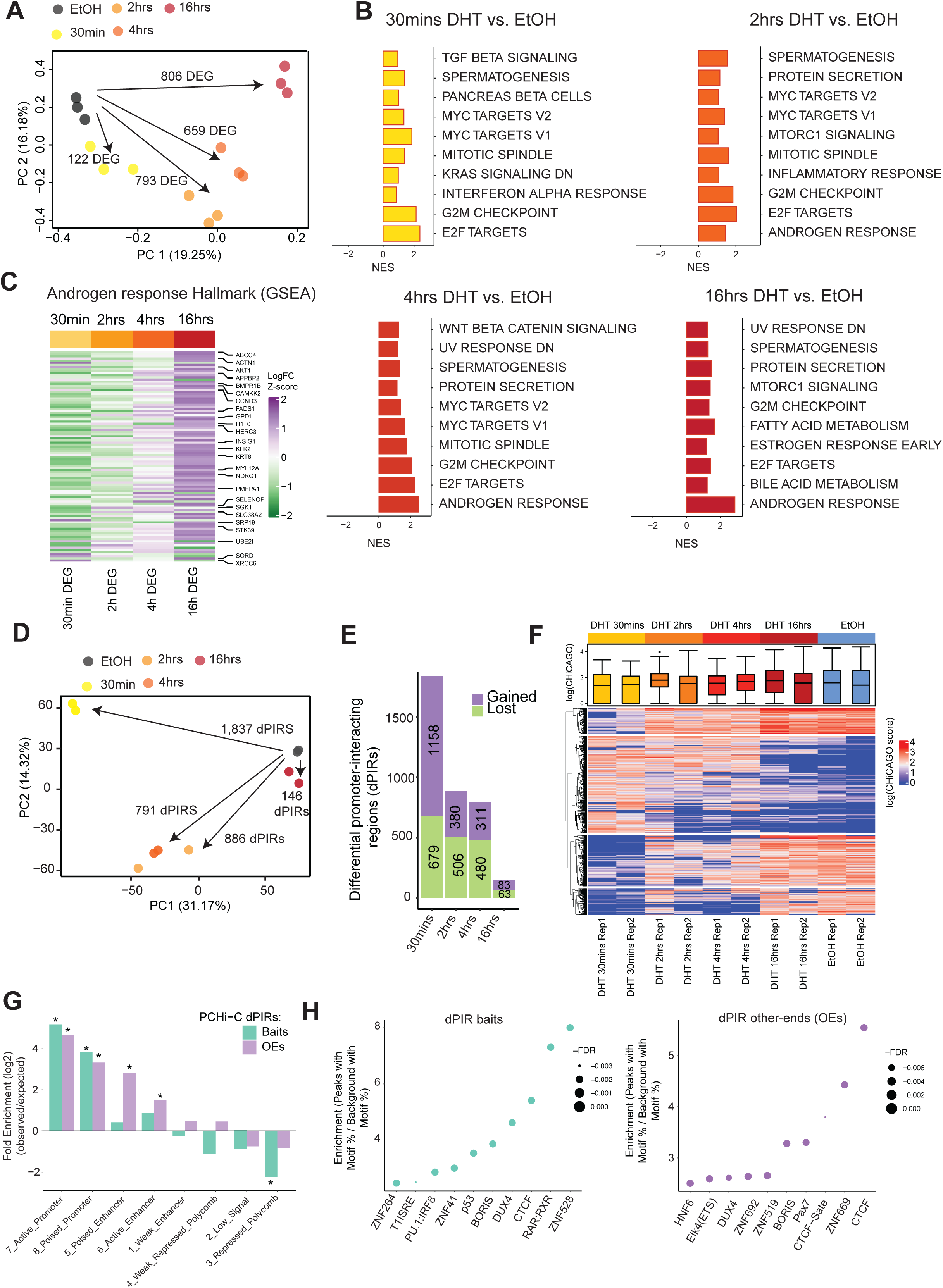
Androgen-stimulation induces progressive changes in AR-driven transcription and transient changes in differential chromatin interactions. **(A)** Principal component analyses showing the relationship among the mRNA transcriptome profiles upon DHT stimulation in the androgen-stimulation time-course. Data plotted for normalised TPM values. The number of differentially expressed genes (DEG) between each DHT time-point and EtOH is shown. **(B)** Normalized enrichment scores (NES) for signature GSEA gene sets representing differentially expressed genes in RNA-seq data at each time-point of DHT stimulation compared to EtOH control. **(C)** Heatmap showing change in expression (Z-scores of logFC vs. EtOH) of genes belonging to the Androgen Response (GSEA) Hallmark. **(D)** Principal component analyses showing the relationship among the chromatin interactions at promoter-interacting regions (PIRs) in Promoter Capture Hi-C data upon DHT stimulation time-course compared to EtOH. Data plotted for top 5000 interactions. The number of differential PIRs (dPIRs) between each DHT time-point and EtOH is shown. **(E)** Number of gained and lost dPIRs detected between each DHT time-point and EtOH control. **(F)** Heatmap of CHiCAGO scores (weighted −log *P* values) for dPIRs detected at each DHT time-point compared to EtOH. **(G)** ChromHMM (LNCaP) annotation (* *P* value < 0.001) of dPIRs compared to matched random regions across the genome. Observed over expected enrichment at dPIR baits and dPIR other-ends (OEs) is plotted. **(H)** Transcription factor motifs enriched at differential chromatin interactions (left dPIRs baits and right dPIRs other-ends) plotted by FDR and percentage of peaks with motif (y-axis) compared to matched random background regions across the DHT time-course.

Gene Set Enrichment Analyses (GSEA)^39,45^ of the upregulated genes identified in the 30-minute DHT treated cells compared with EtOH-treated control identified the top three enriched hallmarks “E2F targets” (FDR q-value < 0.0001, Normalized Enrichment Score (NES) = 2.38), “G2M checkpoint” (FDR q-value < 0.0001, NES = 2.14) and “MYC targets” (p = 4.2 × 10^−4^, NES = 1.86), suggesting that cell-cycle genes and potential oncogenes may be involved in early response to androgen stimulation (**Fig. 2B**). At 2 hours post-DHT stimulation, upregulated genes were enriched for another cell-cycle related hallmark, “mitotic spindle” (FDR q-value = 0.001, NES = 1.61), as well as prostate-specific hallmarks, including “spermatogenesis” (FDR q-value = 0.002, NES = 1.54) and notably the “androgen response” hallmark (FDR q-value = 0.028, NES = 1.44). “Androgen response” genes were similarly enriched at the 4-hour time-point (FDR q-value < 0.0001, NES = 2.48) and at 16-hours (FDR q-value < 0.0001, NES = 2.86) (**Fig. 2B-C, Supp. Fig. 2E**). In contrast, the MYC target gene set (V1 and V2) showed increased expression at 30-minutes, but notably the expression of these genes rapidly reduced after 4 hours of DHT-treatment (**Supp. Fig. 2F**).

Next, to investigate the role of changing 3D chromatin interactions in DHT-induced AR binding, we measured genome-wide promoter-interacting region contacts across four time points following androgen treatment compared with EtOH control cells, using Promoter Capture Hi-C (PCHi-C). PCHi-C data normalization and signal detection using the CHiCAGO pipeline identified on average 31,610 significant cis-interactions between baited promoters and other genomic regions (promoter-interacting regions, PIRs) (**Supplementary Table 3**). We found a strong concordance in pairwise interaction read counts between the biological replicates from the same treatment and a clear separation between time-points of DHT treatment (**Fig. 2D**). Surprisingly, PCA of ChICAGO scores for the top 5,000 chromatin interactions revealed a different trajectory of promoter interaction changes than those for transcription, with hormone-deprived EtOH control cells clustering together with 16-hours DHT-treated cells and early time-points cells (30-minutes, 2-hours and 4-hours) clustering together (**Fig. 2D**).

We next used the calculated ChICAGO interaction scores to identify differential PIRs (dPIRs) using chicdiff ^23^ at a significance cut-off of FDR < 0.05. Across the time course, we identified 3,660 dPIRs (1,932 gained and 1,729 lost) (**Fig. 2E-F) (Supplementary Table 4)**. Notably, the majority of differential chromatin interactions occurred at 30-minutes post-DHT treatment, and of these, 63% (1158/1837) were gained dPIRs compared to EtOH. In contrast, at 2 and 4 hours post DHT treatment, the majority (∼60%) of differential chromatin interactions were lost (2 hours; 506/886 and 4 hours; 480/791) (**Fig. 2E-F**). At 16 hours post-DHT stimulation we identified only 146 dPIRs; 56.8% of which were gained (**Fig. 2E-F**). dPIRs across the time course were also significantly enriched for ChromHMM enhancers and promoters (**Fig. 2G**) and structural proteins (CTCF, BORIS) as well as other zinc-finger transcription factors (**Fig. 2H**).

### Multi-omics analyses identify patterns of temporal variation after androgen stimulation

To provide further insights into the relationship of the different molecular temporal responses leading to a change in AR-induced gene expression, we performed an integrated multi-omics analysis of the temporal sequence of changes, using the statistical framework, MOFA+^41^. This tool identifies temporal patterns of variation across the DHT treatment time-course by applying MEFISTO modelling^42^, which allows for identification of non-temporal and temporal variation in the data and characterization of dataset-specific and shared factors. After filtering AR ChIP-seq, RNA-seq and PCHi-C temporal datasets based on (i) differential AR binding (FDR < 5%, logFC > 1; 20,340 regions), (ii) differential gene expression (FDR < 5%, logFC > 1; 1,921 regions) and (iii) CHiCAGO scores of differential chromatin interactions (FDR < 10% by chicdiff/limma; 1,197 regions) (**Fig. 3A**), we applied the MOFA+ MEFISTO model and identified three distinct patterns (Factors) of temporal variation during DHT treatment relative to the EtOH control in LNCaP cells (minimum explained variance 20% in at least one dataset) (**Fig. 3B**). Cumulatively, the three Factors explained the temporal variation induced by DHT stimulation, including 80.18% in the AR binding data, 65.51% in the gene expression data and 49.58% in the 3D chromatin interaction data (**Fig. 3B**) (**Supplementary Table 5**). The majority (62.21%) of differential AR ChIP-seq binding temporal variance is explained by Factor 1 and Factor 3 (22.61%, **Fig. 3B** and **Supp. Fig. 3A-B**). Factor 2 captures 49.50% temporal variation in the 3D chromatin interactions data, with interactions being largely captured only in Factor 2 (**Fig. 3B** and **Supp. Fig. 3C**). Differential gene expression is partially explained by all 3 f\Factors, with Factor 1 and Factor 2 capturing 19.16% and 16.15%, respectively and Factor 3 capturing 36.85% of the changes (**Fig. 3B** and **Supp. Fig. 3D-F**). The Factors 1 and 2 were largely orthogonal (R^2^ Factor 1 = 0.279, R^2^ Factor 3 = 0.301) and Factors 1 and 3 were correlated (R^2^ = 0.780) (**Fig. 3C**) (**Supplementary Table 6**). Factor 1 captures nonlinear temporal changes mainly driven by AR binding that rapidly occur at 30-minutes and gradually taper off by 16-hours (**Fig. 3D**). Factor 2 captures fast changes driven by 3D chromatin interactions that primarily occur by 30-minutes and are maintained for 4 hours and then recover by 16-hours (**Fig. 3D**). Factor 3 captures linear progressive changes driven mainly by gene expression and AR binding across all time-points (**Fig. 3D**). These three Factors, which provide a framework for the characterization of different patterns of temporal variance in gene expression and chromatin interactions in response to DHT-induction of AR binding, are explained by different subsets of regions from each dataset and will be referred to further in the manuscript by their driving dataset and trend over time, namely “Fast AR binding” (Factor 1), “Transient Chromatin Interactions” (Factor 2) and “Progressive Gene Transcription” (Factor 3).

**Figure 3:**
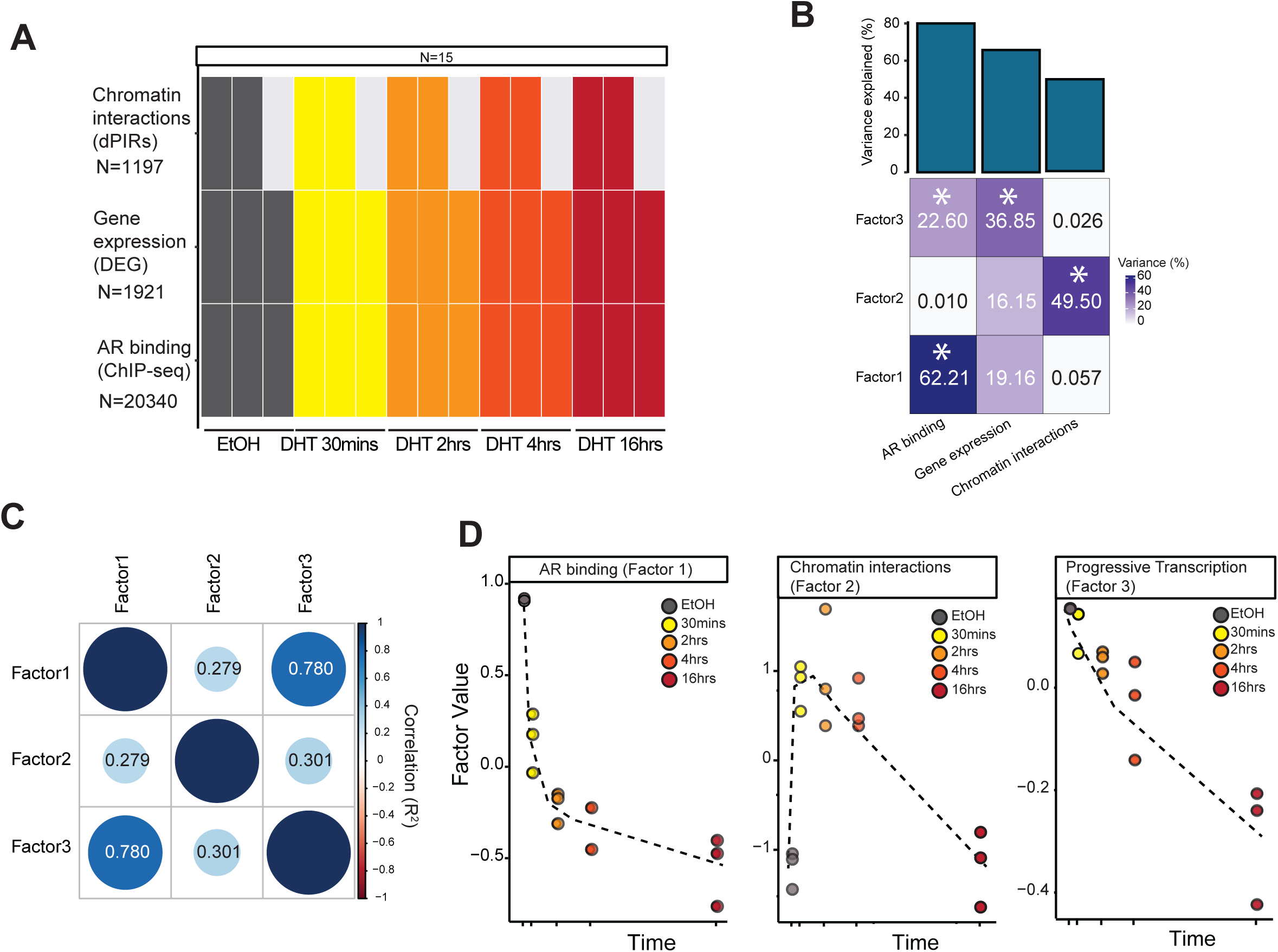
Integrated multiomics analyses of androgen-stimulation in LNCaP cells. **(A)** Illustration of the input data covering chromatin interactions (dPIRs; N = 1,197, replicates = 2), gene expression (DEG; N = 1,921, replicates =3) and AR binding (N = 20,340, replicates = 3) across the DHT time-course and EtOH control). **(B)** Percentage of variance explained by the MOFA+ MEFISTO model. The barplot (top) shows the percentage of variance explained by all factors, and the heatmap (bottom) shows the values for individual factors. The percentage of variance explained is shown. Asterisk (*) indicates significantly contributing datasets for each factor with a minimum of 20% variance explained. **(C)** Pairwise Pearson correlation (R^2^) between the three latent factors identified in **B**. **(D)** Learnt factor values plotted as a function of DHT treatment time for each factor identified. Dots correspond to individual factor values per DHT time-point and EtOH.

### Androgen stimulation results in progressive transcription of canonical AR response genes

The “Progressive Gene Transcription” subset (Factor 3) revealed the temporal relationship of concordant subsets of gained AR binding and differentially expressed genes in response to DHT stimulation, with only a small contribution of chromatin interactions in this subset (**Fig. 3B**). GSEA using the Hallmark gene sets confirmed that “Progressive Transcription” subset is significantly enriched only for the “ANDROGEN RESPONSE” gene set (**Supp Fig. 4A**), including 41 differentially expressed genes (P value = 1.2 x 10^-24^, ChIP-Enrich Effect = 1) (**Fig. 4A** and **Supp. Fig. 4B**). Interestingly these genes progressively increase in differential expression (logFC) over time, with more weight being contributed by genes that become upregulated by 4 and 16-hours (**Fig. 4A**). Additionally, we analysed AR signal at the top weighted AR binding sites in this factor and saw that AR signal increases linearly over time, matching the temporal trajectory of the differential gene expression in this Factor (**Supp Fig. 4C-D**).

**Figure 4:**
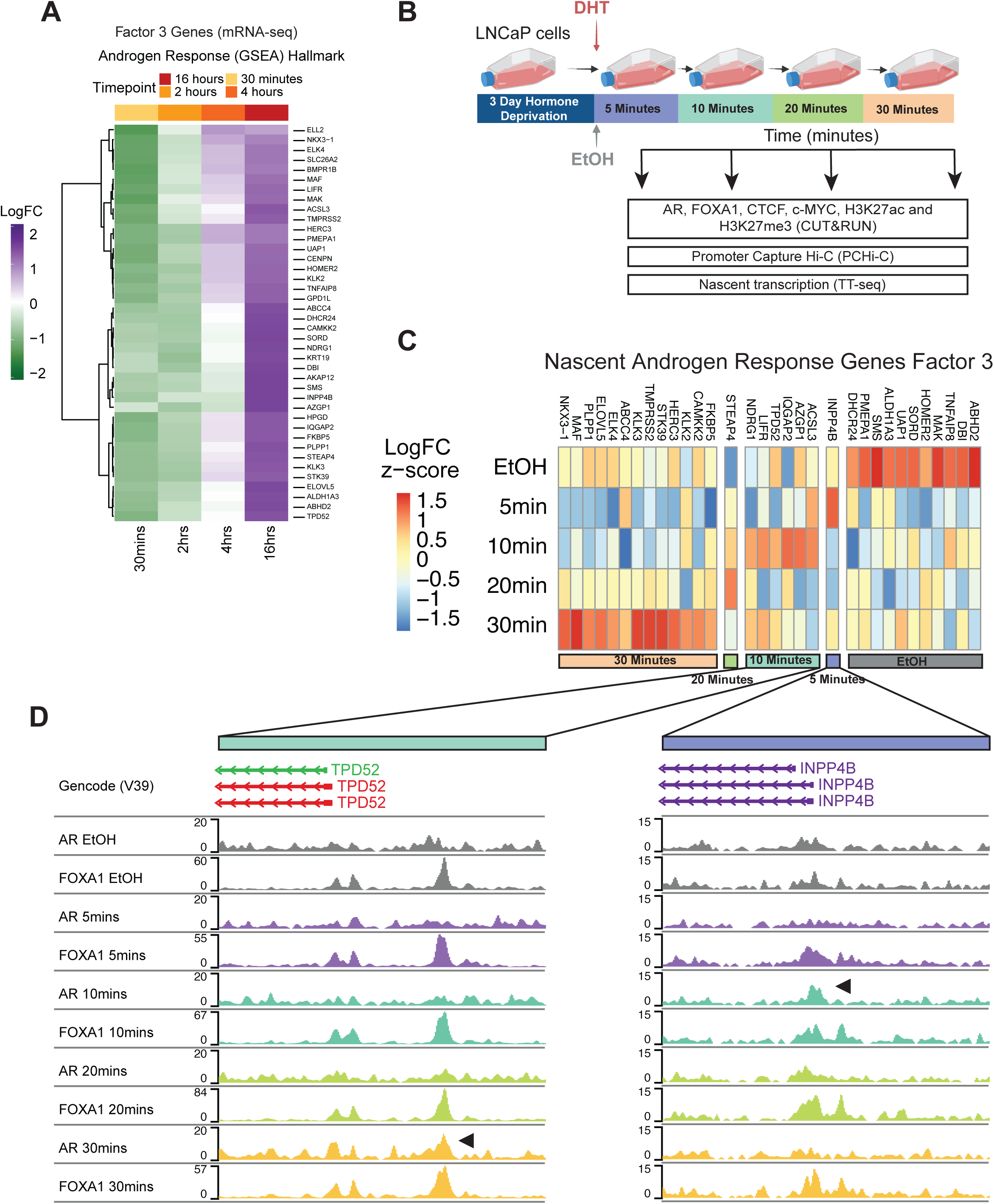
Androgen-stimulation results in progressive transcription at canonical AR response genes. **(A)** Heatmap showing change in expression (Z-scores of logFC vs. EtOH) of genes belonging Androgen Response (GSEA) Hallmark in Factor 3 (“Progressive Transcription”). **(B)** Schematic overview of the experimental design for the “early” DHT time-course. LNCaP cells were treated with 10nM DHT and samples were collected at five time-points for the androgen-stimulation time-course (5mins, 10mins, 20mins, 30mins and EtOH (control) for CUT&RUN (AR, FOXA1, CTCF, c-MYC, H3K27ac, replicates = 3), Promoter Capture Hi-C (PCHi-C, replicates = 2) and TT-seq (replicates = 2). **(C)** Heatmap showing nascent expression (Z-scores of logFC 4sU labelled TT-Seq vs. unlabelled Total RNA-Seq) across “early” DHT treatment time-course of genes in the Androgen Response Hallmark and with top weights in Factor 3 (“Progressive Transcription”). **(D)** Representative examples demonstrating the association between FOXA1 and AR binding as compared across DHT time-points and nascent expression of top weighted androgen response hallmark genes: *TPD52* (left panel) and *INPP4B* (right panel). FOXA1 and AR CUT&RUN signal tracks (replicates merged) are shown. Arrow indicates the initial AR co-binding event with FOXA1 induced by the androgen stimulation.

To next determine if the change in AR binding precedes or occurs simultaneously with AR-mediated gene transcription, we characterized the more immediate temporal impact of AR activation by generating an earlier time-course of androgen stimulation (**Fig. 4B**). We collected DHT-treated LNCaP cells at 5 minutes, 10 minutes, 20 minutes and 30 minutes time-points and characterized transcription factor binding (AR, c-MYC (MYC) and FOXA1 CUT&RUN), histone modifications (H3K27ac CUT&RUN) and chromatin interactions (PCHi-C) at each of early time-points (**Fig. 4B**).

Additionally, we captured the transient rate of androgen-induced gene expression using nascent transcriptome profiling with TT-seq^20^ (Supplementary Table 8). As mRNA-seq used in the MOFA+ model only captures steady-state mRNA transcripts, we first used Total RNA-seq and TT-seq to capture nascent transcription at each of the early time-points (**Supp Fig. 4E**). This revealed that 49% (79 of 161) of the top “Progressive Transcription” genes are already nascently transcribed under the hormone-deprivation conditions with control EtOH treatment and not directly induced by androgen stimulation, unlike the remaining nascent genes that were activated across the time course (**Supp Fig. 4E**). Unlike non-nascent genes we found that all nascently expressed genes were enriched for AR, FOXA1 and H3K27ac, with the highest FOXA1 and AR binding observed at these nascent genes at 20-min after DHT stimulation. Acetylation of H3K27 at the flanking ARE nucleosomes shows maximal enrichment after 5-minutes of DHT treatment (**Supp Fig. 4F**). We observed similar temporal pattern of FOXA1 and H3K27 acetylation enrichment at all genes belonging to the “Progressive Transcription” subset with maximal FOXA1 and H3K27ac enrichment at 20 mins post-treatment (**Supp. Fig. 4G**).

To further confirm whether the early gain of AR and H3K27ac at “Progressive Transcription” genes upon DHT stimulation, could be a general mechanism in androgen dependent prostate cancer, we performed AR and H3K27ac ChIP-seq experiments at the early DHT time-points (5-mins, 10-mins, 20-mins, 30-mins, EtOH), as well as RT-qPCR for gene expression in an additional prostate cancer cell line; LAPC-4. By comparing DHT-induced gene expression changes at key AR target genes, we confirmed a similar response trajectory to DHT stimulation between the LNCaP and LAPC4 cells (see Supplementary Note). We found that AR binding and H3K27ac signal in LAPC4 cells was also enriched at genes in the “Progressive Transcription” subset (**Supp. Fig. 4H**). Similar to LNCaP results the highest enrichment of AR binding was observed at 10-20 minutes after DHT and acetylation of H3K27 showed constitutive enrichment with maximum at 5 minutes of DHT treatment (**Supp. Fig. 4I**), suggesting that this pattern could be a common hallmark of AR-driven prostate cancer.

We next focused on Androgen Response Hallmark genes with top weights in Factor 3 (41 genes). We found that while 11 genes were already nascently expressed in the EtOH control, majority (30 genes) were nascently expressed in response to DHT treatment (**Fig. 4C**). These included known androgen-regulated such as *KLK2*, *KLK3*, *KLK4*, *ELK4*, *TMPRSS2* and *NKX3-1*. The earliest gene being transcribed after DHT stimulation is *INPP4B*, which is nascently transcribed at 5 minutes and differentially expressed in the mRNA-seq data at 16 hours (**Fig. 4C**). Interestingly, androgen induction of this gene depends on FOXA1 co-occupancy and has been implicated in many hormone-dependent cancers^46-49^. At the *INPP4B* gene locus, FOXA1 recruitment co-occurs with nascent expression within 5 minutes of DHT treatment, which precedes transient AR binding at 10 minutes (**Fig. 4D & Supp. Fig. 4J**). The *TPD52* locus also exemplifies this temporal dynamic, where FOXA1 binding is constitutive, and nascent expression occurs at 10 minutes followed AR co-binding at 30 minutes (**Fig. 4D & Supp. Fig. 4J**).

Taken together, the “Progressive Transcription” subset is characterised by canonical features of androgen-response, such as the activation of core androgen response genes. Moreover, nascent RNA transcription at key AR target genes captured in the early time-course indicates that FOXA1 binding and H3K27ac proceeds RNA transcription and AR binding at these early response genes.

### Androgen-stimulation induces transient 3D chromatin interactions at MYC target genes

Based on the large increase in transient 3D chromatin interactions present at 30 minutes of DHT stimulation, we next explored the “Transient Chromatin Interactions” subset (Factor 2). This subset is characterized by a fast differential change in chromatin interactions (variance explained 49.5% by dPIRs and 16.15% by gene expression) and subsequent resolution after the initial DHT treatment. The 3D chromatin interaction changes are largely occurring by 30-minutes and resolving between 4- and 16-hours of DHT stimulation (**Fig. 3D**). To assess the importance of architectural proteins contribution to these transient chromatin interactions, we analysed the enrichment of CTCF binding across DHT-treatments at the Factor 2 regions identified in MOFA+. Factor 2 interactions were most significantly enriched for CTCF/BORIS motifs (**Supp. Fig. 5A**) and ∼29% of dPIRs in Factor 2 (150 out of 516) were located at either CTCF motifs or overlapped a CTCF binding site in ReMap2022. These interactions were enriched for CTCF across all early time-points as well as in EtOH control cells (**Supp. Fig. 5B**).

Considering that majority of chromatin interactions in Factor 2 were transiently observed at 30 minutes of DHT stimulation, we expanded our analyses to PCHi-C data in the early time-course (**Fig. 4B, Supplementary Table 9**). We first investigated which of the Factor 2 chromatin interactions changed between EtOH, 5 minutes, 10 minutes, 20 minutes and 30 minutes of DHT treatment. Initially, we compared the number of PIRs detected at each of the early DHT time-point to Factor 2 chromatin interactions and found that around ∼50% (483 PIRs) of Factor 2 regions overlapped PIRs detected in the early DHT treatment time-course PCHi-C data (**Supp. Fig. 5C**). We next compared the ChICAGO scores at each time-point for PIRs overlapping a Factor 2 region and those that did not overlap. We found a significant (P < 0.05) increase in the ChICAGO scores for chromatin interactions at Factor 2 regions at 20 minutes and 30 minutes after DHT stimulation, while all other PIRs did not exhibit any significant change (P > 0.05) (**Fig. 5A**). Supporting this finding, we also observed a gain in contact frequency at 20 minutes for all PIRs in Factor 2 (**Fig. 5B**). This increase in ChICAGO scores at Factor 2 dPIR regions was maintained at 2 hrs, partially resolved at 4 hrs with majority of contacts largely lost by 16 hrs after DHT stimulation (**Supp. Fig. 5D**).

**Figure 5:**
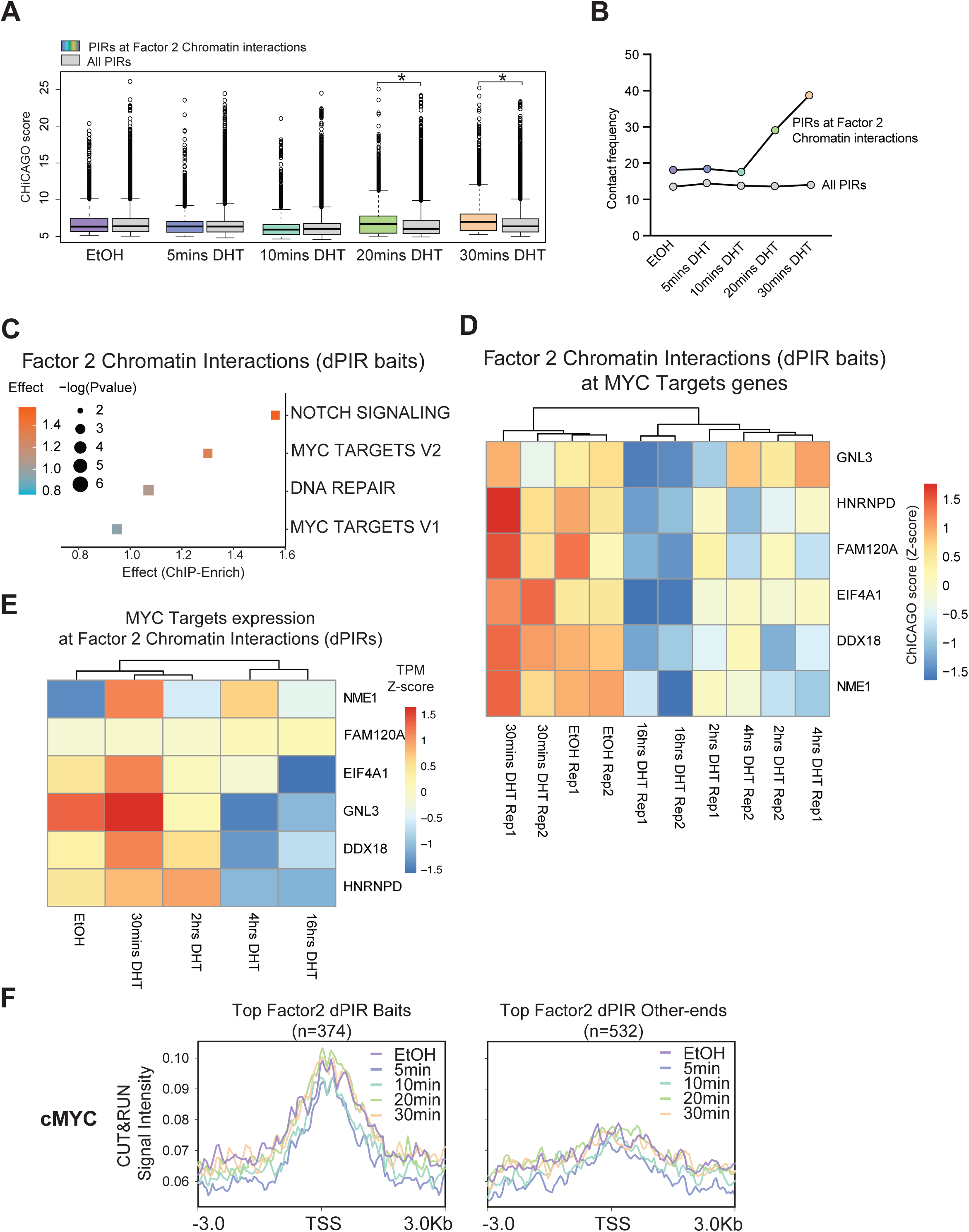
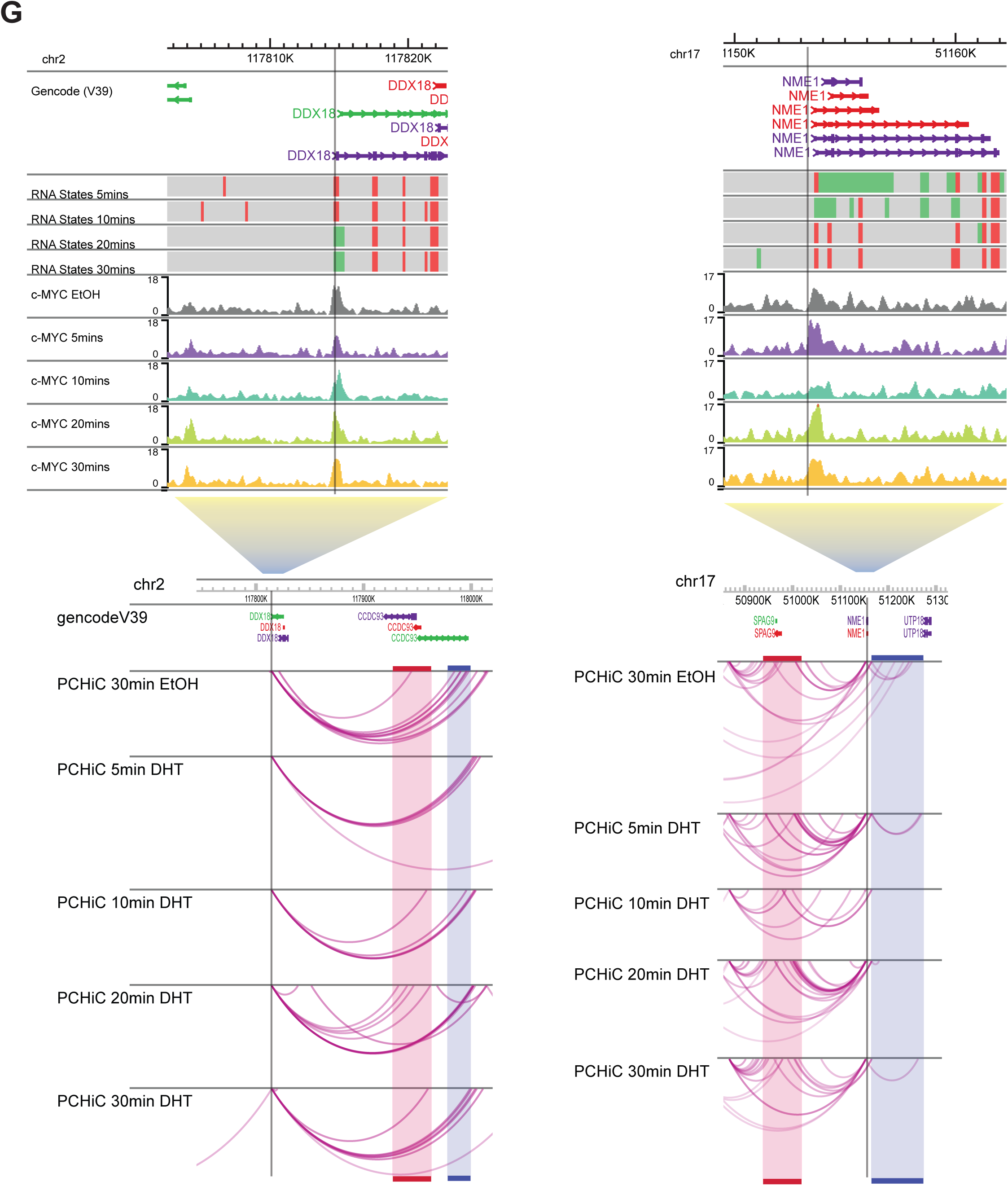
Androgen-stimulation results in transient chromatin interactions. **(A)** Box plot of CHiCAGO scores (weighted −log P values) for Factor 2 (“Chromatin interactions”) PIRs and all PIRs in the early DHT time-course. Asterisk (*) indicates P value < 0.05 (two-sided *t*-test). **(B)** Contact frequency (number of OEs per PIR) for Factor 2 (“Chromatin interactions”) dPIRs and all PIRs in the early DHT time-course. **(C)** Effect scores (ChIP-Enrich) for signature gene sets representing Factor 2 chromatin interactions (dPIRs). Hallmark gene sets with significant enrichment (P value < 0.05, effect score >1) are shown. **(D)** Heatmap of CHiCAGO scores (Z-scores of weighted −log *P* values) for dPIRs belonging to top 10% Factor 2 regions at genes belonging to MYC Targets (V1/V2) (GSEA) Hallmark. **(E)** Heatmap showing change in expression (Z-scores of TPMs) of genes belonging to the MYC Targets (V1/V2) (GSEA) Hallmark that overlap a top 10% Factor 2 dPIR bait region. **(F)** Average CUT&RUN signal intensity for c-MYC binding at dPIR baits (left) and other-ends (right) included in top 10% Factor 2 dPIRs for each DHT treatment time-point. **(G)** Representative examples demonstrating the association between transient 3D chromatin interactions and c-MYC binding to nascently expressed MYC target genes as compared across DHT time-points: *DDX18* (left panel) and *NME1* (right panel) genes. RNA states, c-MYC CUT&RUN signal and PCHi-C data (arc) tracks are shown. Highlighted regions include dPIRs with other-ends that are gained (red) and lost (blue) for each time-point of the early DHT stimulation compared to EtOH control.

To determine the genes involved in the early transient 3D chromatin interactions, we analysed the gene set enrichment of baited promoters involved in these Factor 2 top weighted dPIRs. We found that the differential interactions were significantly enriched for hallmark gene sets (P value < 0.05, effect score > 1) “NOTCH SIGNALLING”, “MYC TARGETS V2” and “DNA REPAIR” (**Fig. 5C**). Notably our data showed an enrichment of 3D chromatin interactions at MYC target genes (**Fig. 5C**), together with an early change from “MYC Targets” (**Supp. Fig. 2F**) to “Androgen Response” gene expression (**Fig. 2C**), following DHT stimulation. MYC has previously been reported as a major driver of prostate cancer tumorigenesis^50^. To further investigate the relationship between MYC targets and Factor 2 top 3D chromatin interactions we further explored the Promoter Capture Hi-C data and found that enrichment for MYC target genes was specifically driven by 226 dPIR regions located at 8 genes belonging to the “MYC TARGETS V2” hallmark, which overlapped six MYC target genes (**Fig. 5E**). Overall, we observed high ChICAGO interaction scores in EtOH and 30-minute DHT samples, which decreased with the DHT treatment time and were the lowest at 16-hours post-DHT (**Fig. 5D**). We also observed high ChICAGO interaction scores in EtOH and 30-minute DHT samples, which decreased with the DHT treatment time and were the lowest at 16-hours post-DHT (**Fig. 5D**). Of these, all six MYC target genes had 3D chromatin interactions that were present in the early DHT treatment (**Supp. Fig. 5E**) and majority showed increased chromatin contacts with the DHT treatment time. We next investigated the mRNA expression of these six MYC target genes and found that high expression of these genes was observed in EtOH control, and in 30-minutes DHT treated samples, with 16-hours DHT samples showing the lowest expression (**Fig. 5E**). Three of these genes *(EIF4A1*, *HNRNPD* and *NME1*) were nascently transcribed as early as 5 minutes after DHT treatment (**Supp. Fig. 5F**), suggesting that transcription of these genes might be partially induced by hormone starvation, rather than DHT treatment (**see Supplementary Note**).

Finally, we explored the MYC binding enrichment at these top Factor 2 dPIRs and found MYC binding at promoter “bait” regions that was maintained across the early DHT time-course, while “other-end” regions did not show any MYC binding (**Fig. 5F**). Visual evidence of the temporal relationship between transient (lost and gained) 3D chromatin interactions and nascent gene transcription at these regions is exemplified at MYC target genes *DDX18* and *NME1* (**Fig. 5G**).

Together, these analyses revealed a dynamic switch in 3D chromatin interactions that occurs between 20 minutes and 4 hours of DHT treatment, with a subset of the fast transient 3D chromatin interaction alterations occurring at MYC target genes.

### Androgen stimulation induces rapid AR binding at constitutive FOXA1 binding sites at novel activated enhancers

The “Fast AR binding” subset (Factor 1) was characterized by rapid temporal changes driven by AR binding (62.2%) and minor contribution of differential gene expression (19.16%) (**Fig. 3B**). Within this subset, approximately half of the temporal variance observed occurs at 30-minute time-point (**Fig. 3D**). This is demonstrated by the difference (compared to EtOH) in average factor value of 0.77, which accounts for 52.79% of the total difference over the 16-hour time course (**Fig. 3D, Supplementary Table 7**). Although the “Fast AR” and “Progressive Transcription” subsets are correlated (R^2^ = 0.78) (**Fig. 3C**), differential AR binding sites with weights in the top 10% of the “Fast AR” subset have an inversely proportional trend when comparing their loading weight against the “Progressive Transcription” weight (**Supp. Fig. 6A**). This suggests that sites in the top 10% will best distinguish the temporal dynamics elucidated by their respective subsets (**Supp. Fig. 6A**).

We next explored specific contribution of TF binding and histone modifications at the top 10% Factor 1 AR binding sites by analysing the enrichment of AR, FOXA1, H3K27ac and c-MYC in the early DHT stimulation time-course (**Fig. 4B**). We observed that AR binding, as expected, was depleted in EtOH (control) and steadily increased within just 5 and 10 minutes post-DHT stimulation, with the highest enrichment observed at 20 minutes of DHT treatment and this enrichment was maintained at 30 minutes after stimulation (**Fig. 6A & Supp. Fig. 6B**). We found a similar induction of FOXA1 binding, which was moderately depleted in EtOH (control) and steadily increased in binding at 5 and 10 minutes post-DHT stimulation, with the highest enrichment observed at 20 minutes of DHT treatment (**Fig. 6B & Supp. Fig. 6C**). This suggests that there is a low level of pre-marked FOXA1 ARE binding sites that can rapidly increase their AR binding after DHT stimulation (**Fig. 6B**). Interestingly, despite androgen depletion, a low level of active histone mark H3K27ac remained present in the “Fast AR” sites in the EtOH control, followed by an increase in the H3K27ac signal observed after DHT stimulation, suggesting a key role in facilitation of nuclear receptor binding at these sites (**Fig. 6C**). Of note, we observed a similar pattern of AR binding induction and H3K27 acetylation in LAPC4 cells, with strong AR depletion in EtOH (control), a rapid increase in AR binding from 5 mins, and maximum enrichment observed at 20 minutes of DHT treatment. Similarity the H3K27ac signal was already present in EtOH and increasing after DHT stimulation (**Supp. Fig. 6D-E**). Moreover, we found that c-MYC binds strongly at some promoters/TSS with top weights in the Factor 1, where AR is also co-localised in LNCaP cells (**Fig. 6D**).

**Figure 6:**
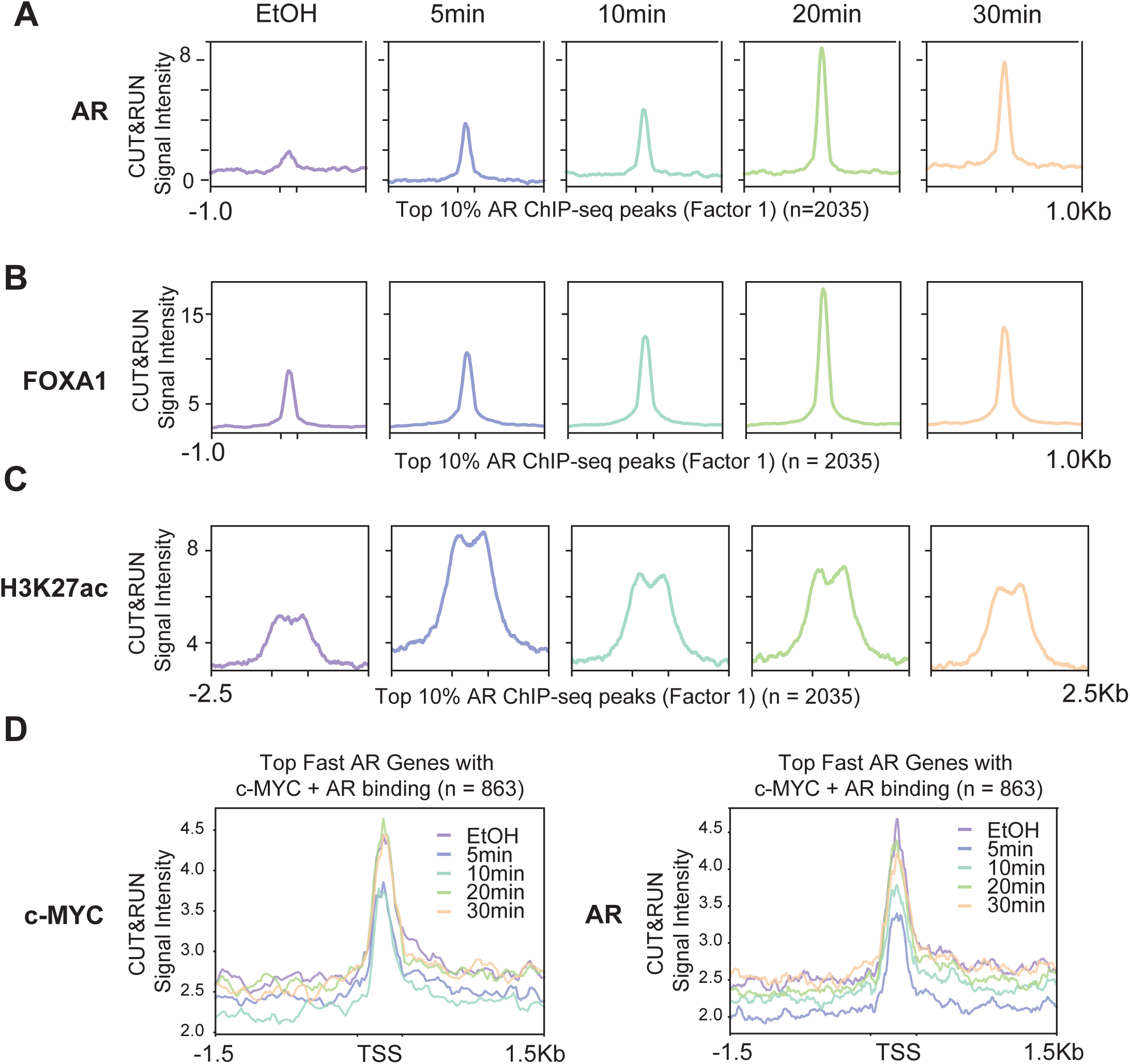
Androgen-stimulation induces rapid AR binding. Average profiles of signal intensity for each time-point of the early DHT stimulation compared to EtOH control; **(A)** AR CUT&RUN; **(B)** FOXA1 CUT&RUN and **(C)** H3K27ac CUT&RUN at top 10% AR ChIP-seq peaks in Factor 1 (“Fast AR binding”); **(D)** c-MYC (left panel) and AR (right panel) CUT&RUN signal intensity at TSS of top differentially expressed genes with c-MYC and AR co-binding in Factor 1 (“Fast AR binding”).

To further elucidate the temporal relationship between this rapid AR and co-regulator binding and resulting gene transcription, we performed GSEA for the nearest genes to AR binding sites. Strikingly, we found that “Fast AR” sites are significantly enriched proximal to the genes belonging to the Androgen Response Hallmark (**Fig. 7A**), which were also identified in the Factor 3 “Progressive Transcription” subset (**Supp. Fig. 3A**). This supports a clear temporal dynamic, where AR is rapidly bound at gene promoters, immediately prior to the initiation of nascent transcription of canonical androgen-responsive genes. For example, this temporal dynamic can be demonstrated at the kallikrein gene locus, where three top-weighted “Fast AR” sites are found proximal to two top-weighted “Progressive Transcription” genes (*KLK2* and *KLK3*) (**Fig. 7B**). Nascent transcription occurs at the “Fast AR” sites, where the most upstream genes appear to be nascently transcribed first. Additionally, the previously described temporal pattern in AR, FOXA1 binding, and H3K27ac enrichment (**Fig. 6A-C**) is recapitulated in this example region (**Fig. 7B**), with similar distribution of AR binding and H3K27ac enrichment observed in LAPC4 cells (**Supp. Fig. 7A**).

**Figure 7:**
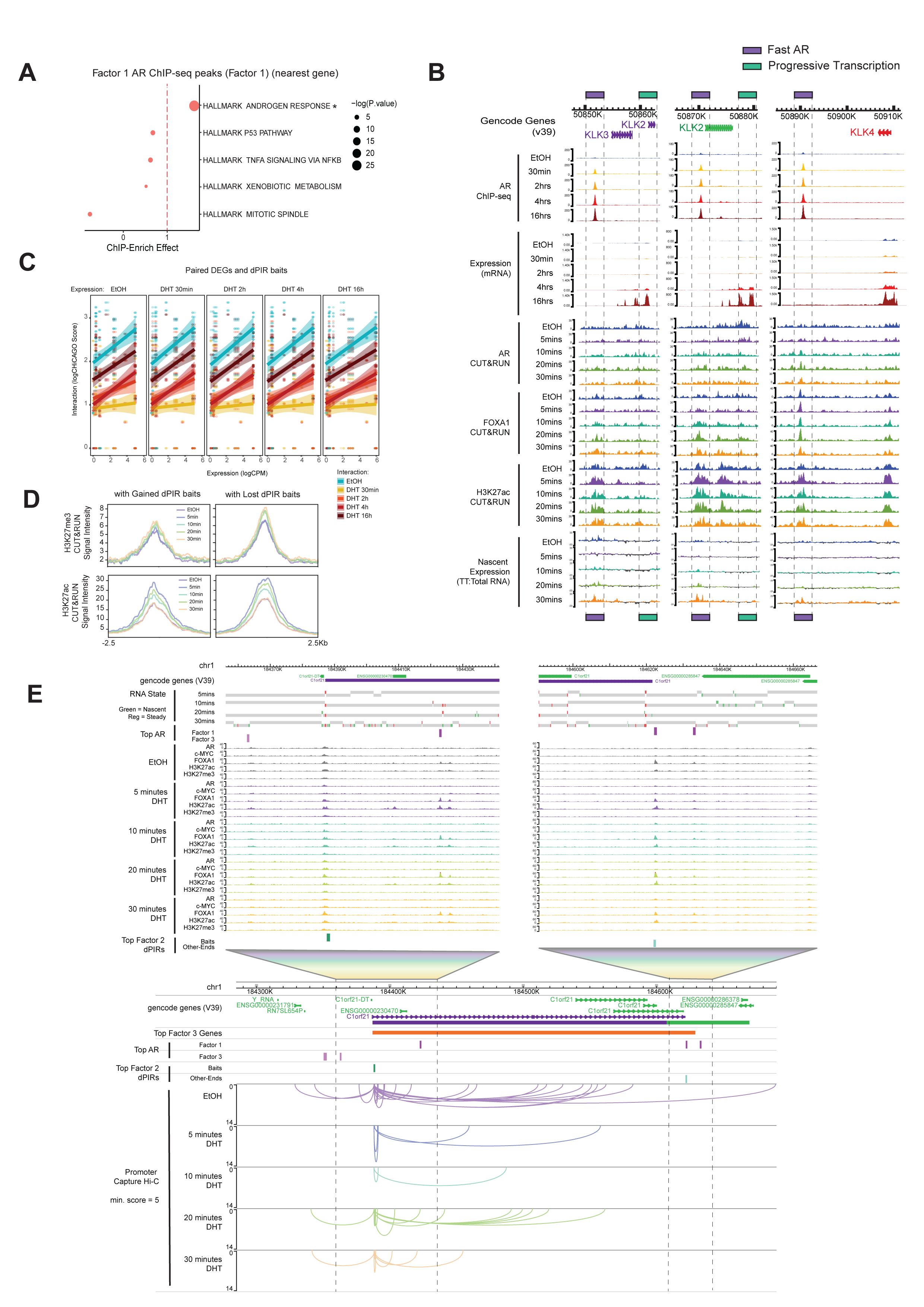
H3K27ac followed by pioneer and transcription factor binding precedes chromatin interaction and transcriptional responses upon androgen stimulation. **(A)** Effect scores (ChIP-Enrich) for signature gene sets representing top 10% AR ChIP-seq peaks in Factor 1 (“Fast AR binding”). * P value < 0.05, effect score >1. **(B)** Representative examples demonstrating the association between “Fast AR” binding sites (purple dashed boxes) and “Progressive Transcription” gene expression (green dashed boxes) as compared across DHT time-points. *KLK* locus (*KLK3*, *KLK2* and *KLK4*) shown. AR ChIP-Seq (n = 3), mRNA-Seq (n = 3), AR, FOXA1 and H3K27ac CUT&RUN (n = 3) and nascent expression (TT:Total RNA-Seq log2 ratio, n = 2) tracks are shown. **(C)** Dot and line plot of interaction score (log(ChICAGO) vs. gene expression (logCPM fold change) of DEGs that have dPIR baits with top weights in Factor 2 (“Transient Chromatin Interaction”) for each time-point of DHT stimulation compared to EtOH control. **(D)** Average profiles of H3K27ac and H3K27me3 CUT&RUN signal intensity at genes which become upregulated between 30 minutes and 2 hours and have dPIR baits with top weights in Factor 2 (“Transient Chromatin Interaction”) that are gained (left panel) and lost (right panel) for each time-point of the early DHT stimulation compared to EtOH control. **(E)** Representative example demonstrating the association “Fast” and “Progressive” AR binding sites (pink track) and “Progressive Transcription” gene expression (orange track) and “Transient Chromatin Interactions” (dPIR baits and OEs, teal track) as compared across the early DHT time-points. *C1orf21* gene loci shown. Nascent RNA state, AR, c-MYC, FOXA1, H3K27ac and H3K27me3 CUT&RUN and PCHi-C data (flat arc, zoomed-out 500 kb region) tracks are shown.

Overall, these results demonstrate that rapid AR binding occurs at canonical androgen response genes and is pre-marked by constitutive acetylation of H3K27 and FOXA1 occupancy, which potentially facilitates the rapid binding of AR.

### H3K27ac and nascent transcription precedes 3D chromatin interaction upon androgen stimulation

To further explore the contribution of androgen-induced 3D chromatin interactions to gene expression in the context of Androgen Response genes, we studied the relationship between 3D chromatin interaction score (dPIRs from CHiCAGO) and differential gene expression (DEG from mRNA-seq) (**Fig. 7C**). We found a strong positive correlation between chromatin interaction score and gene expression in the EtOH control; this correlation is temporally lost at 30 minutes after DHT treatment and re-established from 2 hours and maintained until 16 hours of DHT treatment (**Fig. 7C**).

Due to this loss of correlation, we investigated the chromatin state (H3K27ac and H3K27me3) of TSS with dPIRs for genes that are upregulated between 30 minutes and 2 hours. These analyses revealed an inversely proportional enrichment of H3K27 acetylation and trimethylation at dPIR baits (**Fig. 7D**). At dPIR baits that are gained with DHT stimulation, we found that there was a rapid temporal increase in H3K27 acetylation, with the highest enrichment at 5 mins post-DHT (**Fig. 7D**). Inversely, we observed a moderate enrichment in H3K27me3 at these gained dPIR baits at 30 mins post-DHT treatment (**Fig. 7D**). We also observed a similar trend in chromatin states at lost dPIR baits that show a temporal decrease in 3D chromatin contacts (**Fig. 7D**).

The temporal order is visually exemplified at the *C1orf21* gene loci, where transient decoupling of gene expression (Factor 3) and 3D chromatin contacts (Factor 2) is observed concordant with AR (Factor 1) and co-factor binding, gain of active of histone modifications and initiation (*C1orf21*) of nascent transcription (**Fig. 7E**). Whereas at the *EPHA4* gene loci, we observe a similar transient decoupling but now in association with transcriptional repression (**Supp. Fig. 7B**). Here, gain in repressive histone modifications and transient 3D chromatin contacts (Factor 2), that are subsequently bound by progressive AR (Factor 3), is concordant with repression of this fast AR gene (Factor 1).

Together, our results demonstrate that early changes in chromatin state precede transcription factor binding and transient 3D chromatin interactions which contribute to the regulation of androgen-induced nascent transcription.

### CRISPR-mediated inhibition confirms functional role of rapid and progressive AR sites in AR-mediated gene expression

To next experimentally validate the functional role of key MOFA+ identified AR sites, we tested the effects of ablating AR binding on gene expression alterations at key androgen response genes. Using the CRISPR inhibition (CRISPRi) system (**Fig. 8A**), we targeted a subset of both rapid and progressive AR binding sites that are in close 3D proximity to previously identified AR target genes within the kallikrein gene locus (**Fig. 8B**). These include two “Fast AR” AR binding sites between *KLK15* and *KLK3* genes (ARBS #1 and ARBS #2), a “Progressive Transcription” AR binding site upstream of the *KLK2* gene promoter (ARBS #3), the *KLK2* gene promoter as a positive control (*KLK2* promoter), and a “Fast AR” AR binding site within the *KLKP1* gene (ARBS #4) (**Fig. 8B**). Across the whole kallikrein gene locus, we observed that inhibition of each of the AR binding sites resulted in weakened activation of nearly all genes within the locus, while expression of a known AR target genes outside of this locus (*ZBTB16*) was induced as expected (**Fig. 8C** and **Supp. Fig. 8A**). Importantly, we found that the highest inhibition of *KLK2* gene expression was induced by targeting “Fast AR” (ARBS #2) and “Progressive Transcription” AR binding site (ARBS #3), in addition to the strong inhibitory effect of targeting the *KLK2* promoter directly, as expected (**Fig. 8C** and **Supp. Fig. 8A**). For *KLK4* gene, a significant reduction in gene activation was induced by targeting the “Fast AR” ARBS #2, while *KLKP1* gene expression was most affected by inhibiting “Fast AR” ARBS #4 (**Fig. 8C** and **Supp. Fig. 8A**). DHT induction of *KLK3* gene, which encodes for the Prostate-Specific Antigen (PSA) protein, was significantly reduced by targeting all AR binding sites across the locus, with the strongest effect observed via inhibition of AR binding at either of the ARBS which flank the gene; ARBS #2 (“Fast AR”) and ARBS #3 (“Progressive Transcription”) (**Fig. 8C** and **Supp. Fig. 8A**). Additionally, inhibition of either ARBS #2 or ARBS #3 resulted in reduced *KLK3* encoded PSA protein expression, with the strongest effect observed for “Fast AR” ARBS #2 (**Supp. Fig. 8B & C**). Across the whole kallikrein locus, the strongest inhibitory effects were observed at AR binding sites that had the strongest AR binding and were pre-marked by H3K27 acetylation on their flanking nucleosomes (ARBS #2 and #3 (**Fig. 8B**)).

**Figure 8:**
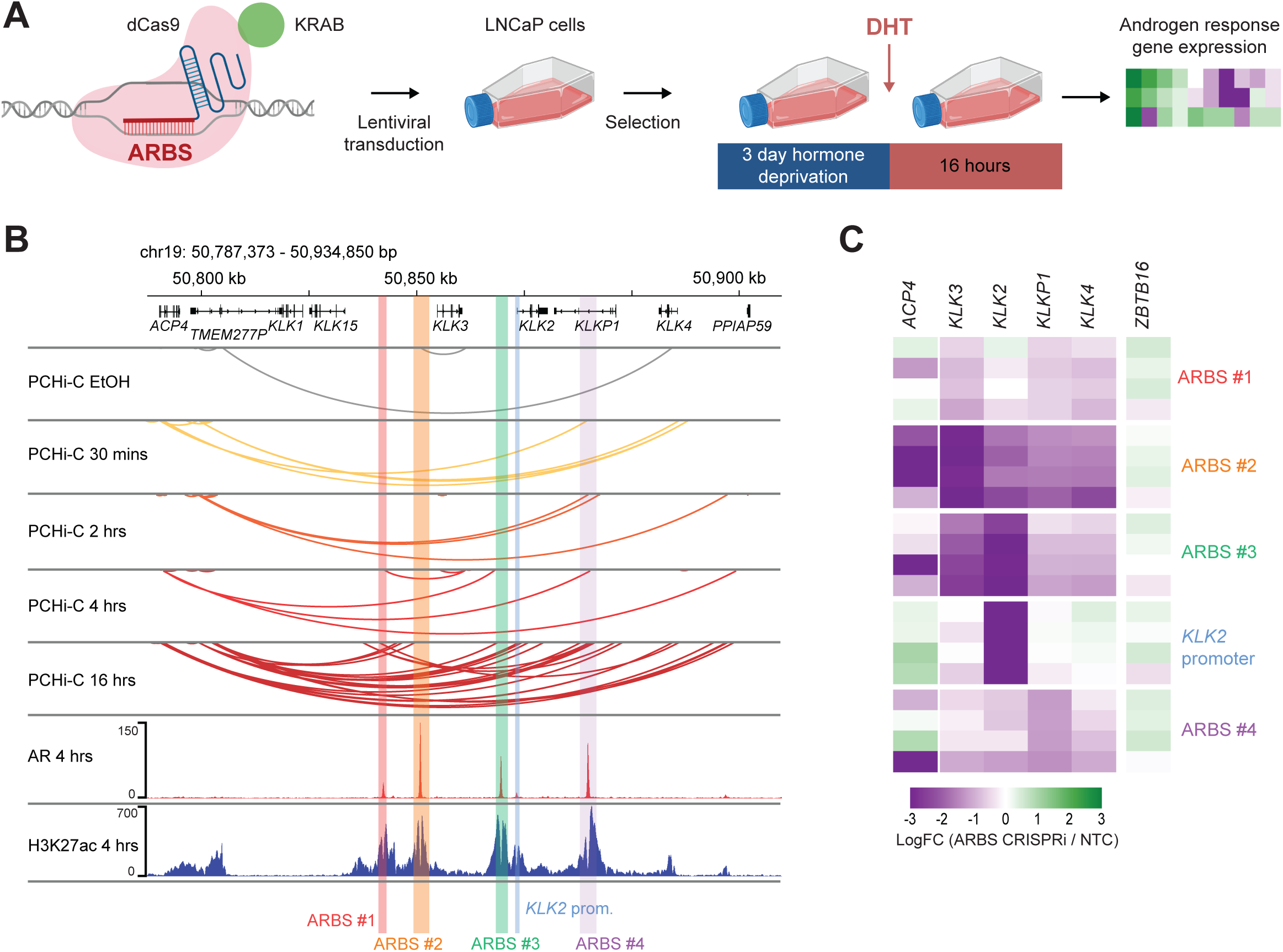
Androgen-mediated transcription is significantly reduced by inhibiting AR-bound enhancer regions. **(A)** Schematic illustration depicting CRISPR experiment of dCas9-KRAB-gRNAs targeting AR binding sites at *KLK* locus in LNCaP cells followed by DHT stimulation **(B)** Genome browser screenshot of the *KLK* gene locus demonstrating guide design for targeting AR binding sites across the locus. AR ChIP-seq (4-hours post-DHT), H3K27ac ChIP-seq (4-hours post-DHT) and PCHi-C across the DHT time-course shown. **(C)** Heatmap showing change in gene expression in CRISPRi compared with CRISPRi-empty controls (NTC) for genes located within the *KLK* locus. Coloured highlights demonstrate the location of AR binding sites and KLK2 gene promoter targeted by CRISPRi.

Finally, we focused specifically on AR binding sites involved in DHT-regulated 3D chromatin interactions within the kallikrein locus. We found that inhibition of AR binding site ARBS #2, which interacts with distal gene *ACP4* via 3D chromatin interactions (**Fig. 8B**) resulted in reduced up-regulation of this gene with DHT treatment (**Fig. 8C** and **Supp. Fig. 8A**). Similar effects were observed for ARBS #4 (**Fig. 8C** and **Supp. Fig. 8A**), which also shows long-range interactions with *ACP4* (**Fig. 8B**). Overall, these results suggest that not all AR binding at AREs contributes equally to DHT-induced gene expression and highlights the importance of both 3D genome and epigenome to spatiotemporal control of transcription.

## DISCUSSION

Here, we report the temporal order of epigenetic-based processes and 3D chromatin interactions associated with androgen induced AR-driven gene expression. We investigated the intricate coordination of gene expression, nascent transcription, 3D chromatin architecture, and transcription factor binding in dynamic models of androgen stimulation in prostate cancer LNCaP and LAPC4 cells. Based on an integrated multi-omics analysis of the temporal sequence of changes, using the statistical framework, MOFA+, we identify three different sequential modes of coordinated interplay of transcription factors, histone modifications, and 3D chromatin interaction dynamics involved in the regulation of androgen response genes (**Fig. 9**).

**Figure 9:**
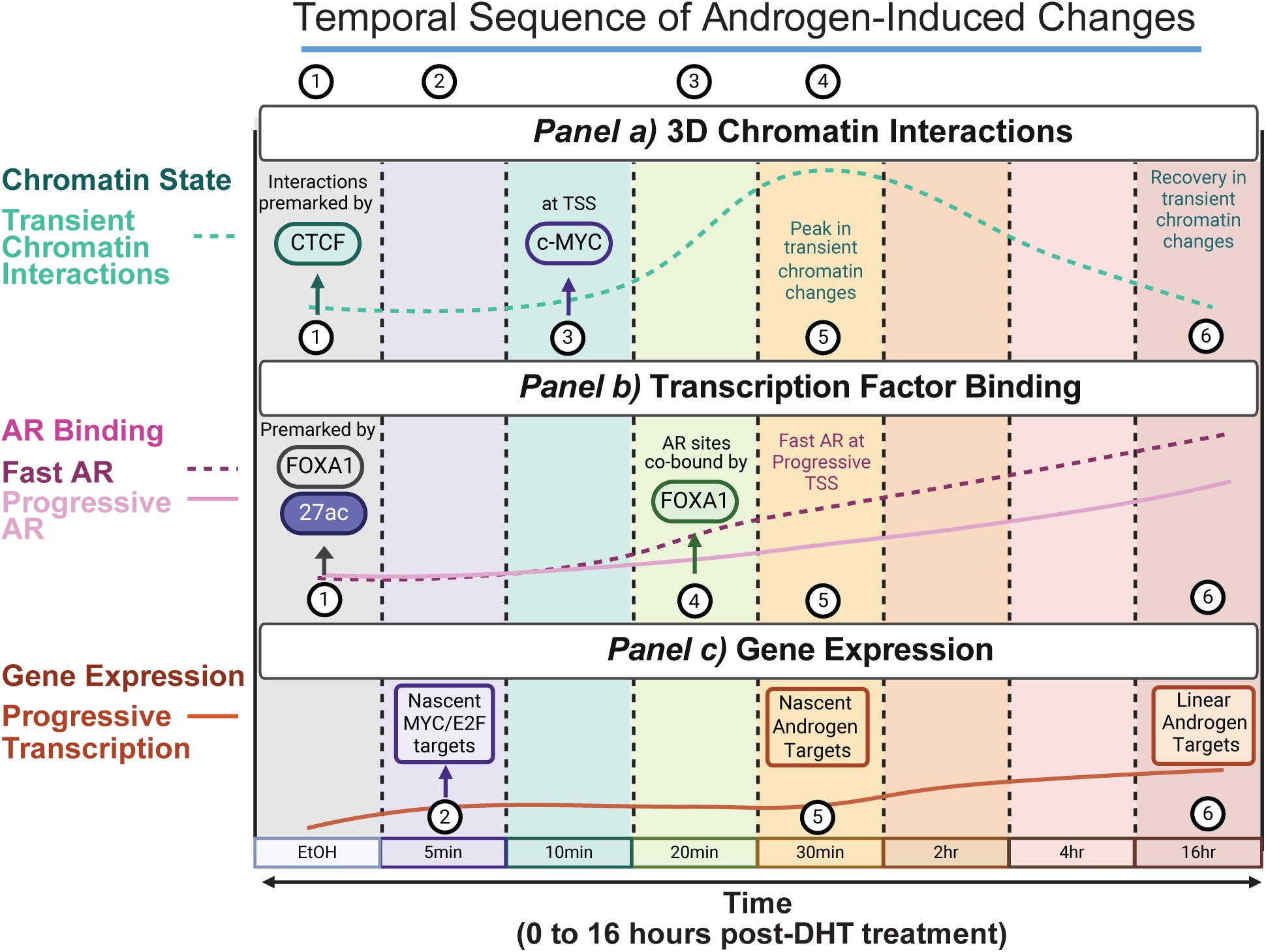
Proposed model of temporal AR-mediated transcription. Model of temporal androgen-induced changes in hormone-responsive prostate cancer, encompassing transcription factor binding, histone modifications, gene expression and 3D chromatin interactions in time. **(1)** Androgen responsive 3D chromatin interactions are pre-marked by CTCF (*Panel a*); rapidly gained AR binding sites are pre-marked by FOXA1 (*Panel b*). **(2)** In the initial 5 minutes, H3K27 is acetylated at transient 3D chromatin interactions (*Panel a*) and at rapidly bound AR sites (*Panel b*); MYC target genes are nascently expressed (*Panel c*) **(3)** At 10 minutes of androgen stimulation, c-MYC is binding at MYC target genes involved in transient chromatin interactions (*Panel a*). **(4)** Within 20 minutes of androgen stimulation, AR and FOXA1 co-bind at “Fast AR” binding sites that are enriched at canonical androgen response genes (*Panel b*). **(5)** After 30 minutes, androgen response genes bound by “Fast AR” (*Panel b)* are nascently expressed (*Panel c*) and transient 3D chromatin interactions show a peak in contact frequency (*Panel a*). **(6)** AR binding is maintained (*Panel b*); transient 3D chromatin interactions are resolved (*Panel a*) and “Progressive Transcription” of canonical androgen response genes linearly increase in expression until the 16-hour time-point (*Panel c*).

First, our temporal model identified a subset of CTCF pre-marked 3D chromatin interactions that were transiently initiated at 20-mins post DHT treatment but were subsequently resolved at later time-points (**Fig. 9; Panel a**). These dynamic 3D chromatin interactions were preceded by nascent expression of MYC target genes at 5 minutes, followed by increased c-MYC binding at their gene promoters at 10 minutes. Recent studies have suggested an antagonistic relationship between expression of canonical androgen response genes and MYC targets^51-53^. However, by extending our study to include earlier time-points after androgen stimulation, we have now shown that 3D chromatin interactions are present at MYC target genes prior to stimulation, and remain up to 30 minutes aftertreatment, followed by a gradual loss of the 3D chromatin interactions. Notably MYC target genes at these transient 3D chromatin interactions are frequently activated in the first 30-minutes of DHT stimulation, followed by decrease in their mRNA expression, which occurs coincidentally with increased expression of androgen response genes at the later time-points. Therefore, our data suggests that MYC is involved in driving the initial transcriptional response to androgen deprivation, prior to the switch to androgen transcriptional response.

Second, our temporal model shows that androgen stimulation induces rapid AR binding at at FOXA1 pre-marked regulatory regions with constitutive H3K27ac (**Fig. 9; Panel b**). While the involvement of AR co-regulators in AR-mediated transcription in prostate cancer have been well-documented, including a suggested role of p300 in inducing H3K27ac at AREs^3,54,55^, the precise timing of initiating steps has not been addressed due to their rapid nature. By including very early time-points in our model, we showed different temporal patterns of gain in AR binding across the ARE sites, which are frequently preceded by modified chromatin states. Similar hormone-induction approaches have previously demonstrated a distinct impact of pioneer transcription factors, such as FOXA1 on gene expression^6,56^. In agreement, our results demonstrate a pioneer role of FOXA1 at AR binding sites gained upon hormone stimulation. We found that a basal level of FOXA1 binding is maintained during hormone starvation followed by a temporal increase in FOXA1 binding strength with androgen treatment over time; highest enrichment of AR/FOXA1 co-binding occurred at 20mins post-DHT stimulation. These FOXA1 bound sites were also enriched in active enhancer-associated H3K27ac marks, before a gain in AR binding and DHT induced nascent transcription of androgen response genes. Although previous studies^57^ suggest AR binding as a trigger for the temporal cascade of AR response, our data shows that AR binding proceeds the initial epigenetic reprogramming of these sites; this difference is likely due to the temporal resolution of our experimental model. Together, our data provides a new temporal dimension to previous studies and supports the concept that enhancer activation is an initial step required for TF binding and chromatin interactions.

Finally, we characterized the contribution of nascent transcription and 3D chromatin interactions to the establishment of AR-mediated transcriptional programs (**Fig. 8; Panel c**). Earlier studies proposed that nuclear receptor binding significantly reorganizes 3D chromatin interactions by recruiting chromatin remodelling complexes or by altering DNA methylation at regulatory regions^58-60^. However, recent work has suggested that gene expression may occur through a more complex change in contact frequency and proximity at already-existing interactions to induce GR- and ER-mediated transcription^16,61,62^. We found that at the later time-points 3D chromatin connectivity was significantly correlated with gene expression, in agreement with previous studies in a similar model of androgen stimulation in LNCaP cells^57^. However, our time-resolved analyses have demonstrated a temporal decoupling of gene expression from contact frequency. During the initial 5-20 minutes of response to stimuli, 3D chromatin interactions are present at MYC target genes and may facilitate their expression, while at the later time-points 3D contacts were present at canonical AR target genes. These observations support our model where we propose that early transcriptional activation leads to transient disruption of 3D chromatin interactions.

Overall, we provide new insights into early contribution of 3D chromatin interactions, TF binding and epigenome modifications to AR-mediated transcription. Due to the complexity of capturing these dynamic changes across a tightly resolved time course, we used two well-characterised models of androgen-dependent prostate cancer, the LNCaP and LAPC4 cell lines. Next, we chose to study TF binding and histone modifications using a recently developed CUT&RUN technology with light cross-linking to preserve the chromatin states at early time-points. Further, we used MEFISTO framework to integrate these temporal multi-omics datasets, enabling us to examine the contribution of multiple regulatory layers at specific genomic regions. This unsupervised approach allowed us more comprehensively dissect how each of the different chromatin and transcriptional features coordinate androgen-mediated transcription, revealing a consistent regulatory model across two independent prostate cancer cell lines. Specifically, we show that androgen stimulation triggers AR binding to pre-marked FOXA1 binding sites and nascent transcription of canonical androgen response genes. Further, we connect 3D chromatin interactions to c-MYC binding and nascent transcription of MYC target genes. Rapid change in 3D chromatin interactions at 30-minutes post-hormone stimulation temporally decouples the relationship between 3D chromatin contacts and gene expression. Importantly, by capturing changes in the first 5-30mins after androgen-stimulation, we revealed that H3K27 nucleosomes are acetylated flanking FOXA1 binding before AR co-binding at these sites, which is interdependent on nascent transcription of the AR target genes. Together our study underscores the importance of high-resolution temporal integrated analysis of multi-omics data to uncover the dynamic epigenetic changes that accompany AR-mediated transcription in prostate cancer.

## ACKNOWLEDGEMENTS

We would like to thank Irina Voineagu and Nicole Green for providing advice and protocols for TT-seq, Daisy Kavanagh, Brian Gloss and the Clark Laboratory members for critical discussions. We thank Arima Genomics for advice on the design of the Promoter Capture Hi-C and National Computer Infrastructure (NCI) for research computing. We thank the South Australian Genomics Centre (SAGC) for next-generation sequencing. The SAGC is supported by the National Collaborative Research Infrastructure Strategy (NCRIS) via Bioplatforms Australia and by the SAGC partner institutes. Graphical Abstract & Figures 1A, 4B and 9 were created with Biorender.com.

## AUTHOR CONTRIBUTIONS

Conception: E.C., F.V.M, S.J.C. & J.A-K.; Design: E.C., A.K., F.V.M, S.J.C. and J.A-K.; Experimental: E.C., G.L-L., G.S., Y.C-S., D.M., T.A.H., C.S., A.J., D.C, A.K., F.V.M. & J.A-K.; Data analyses: E.C., G.L.L., D.V.T, T.J.P, Q.D, K.A.G. & J.A-K.; Writing the manuscript: E.C., S.J.C. and J.A-K.

## CONFLICT OF INTEREST

None declared.

## FUNDING

National Health and Medical Research Council of Australia (NHMRC) Ideas Grants (grant numbers: #1144574, #2028684: #2020334); NHMRC Fellowship and Investigator [grant numbers: #1063559 & #2026430 to S.J.C., #1177792 to Q.D.); The Sylvia and Charles Viertel Senior Medical Research Fellowship (grant number: #SMRF25-10 to J.A-K.); Prostate Cancer Foundation of Australia [grant number: YI-0922 to J.A-K.]; National Computational Merit Allocation Scheme [grant numbers: 2025-63 & 2026-183]. The contents of the published material are solely the responsibility of the administering institution and individual authors and do not reflect the views of the NHMRC. The funders had no role in study design, data collection and analysis, decision to publish or preparation of the manuscript. Funding for open access charge: NHMRC.

## DATA AVAILABILITY

All raw and processed sequencing data generated in this study have been submitted to the NCBI Gene Expression Omnibus (GEO; https://www.ncbi.nlm.nih.gov/geo/) under accession number GSE283232 and GSE333286. The public database of the hg38/GRCh38 genome and annotation files are available from the GENCODE portal (https://www.gencodegenes.org). Biological material used in this study can be obtained from the authors upon request.

## SUPPLEMENTARY FIGURE LEGENDS

**Supplementary Figure 1:**
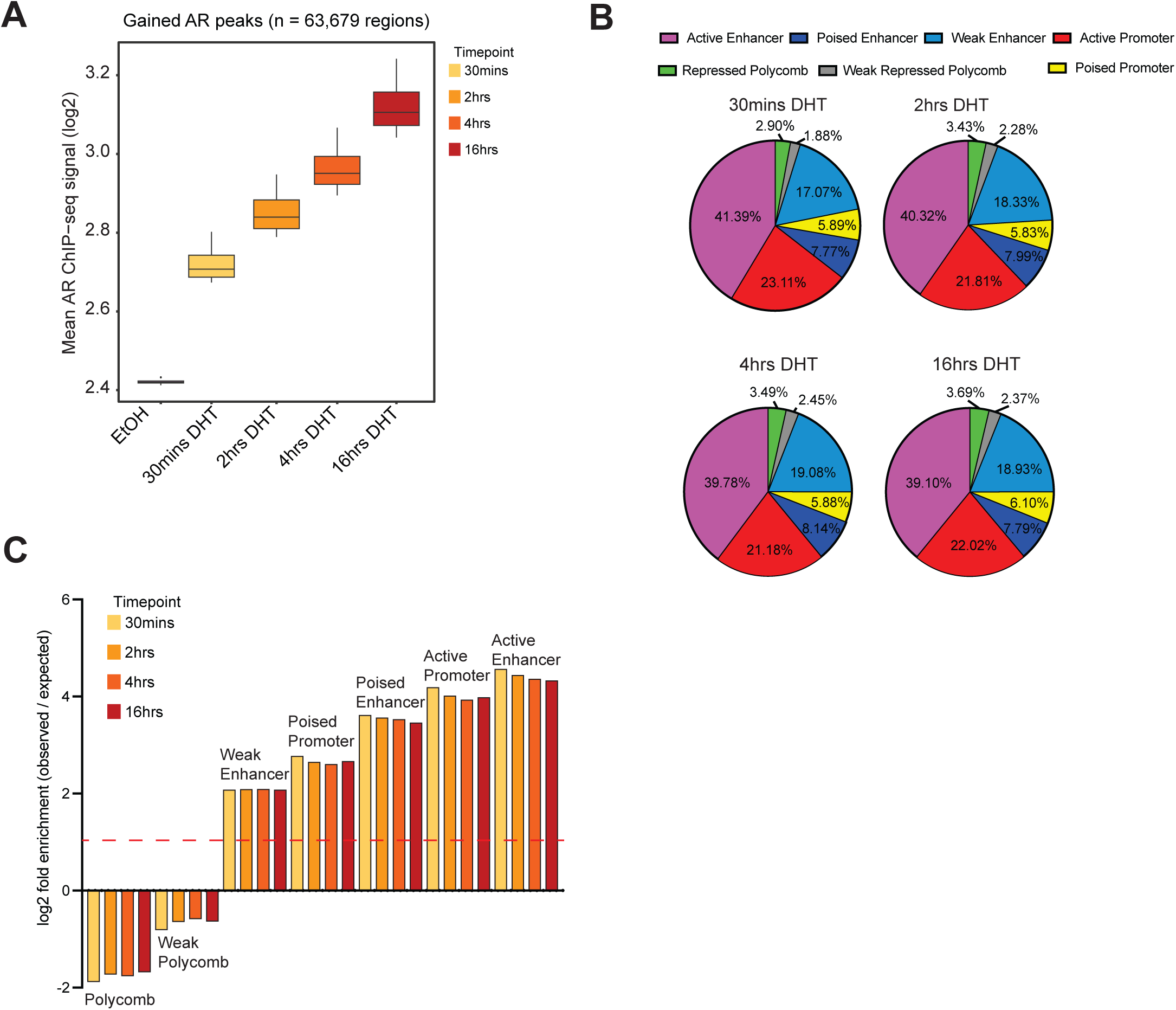
Androgen-stimulation induces changes in AR binding, AR-driven transcription and chromatin interactions. **(A)** Box plot quantification of AR ChIP-seq average signal profile at gained AR binding sites. **(B)** Differential AR ChIP-seq peaks occupancy at ChromHMM regions in LNCaP. The proportions of ChromHMM states distributed at each set of differentially gained AR binding sites. **(C)** ChromHMM (LNCaP) annotation (* *P* value < 0.001, permutation test) of differentially gained AR binding sites per DHT time-point compared to matched random regions across the genome. Observed over expected enrichment is plotted.

**Supplementary Figure 2:**
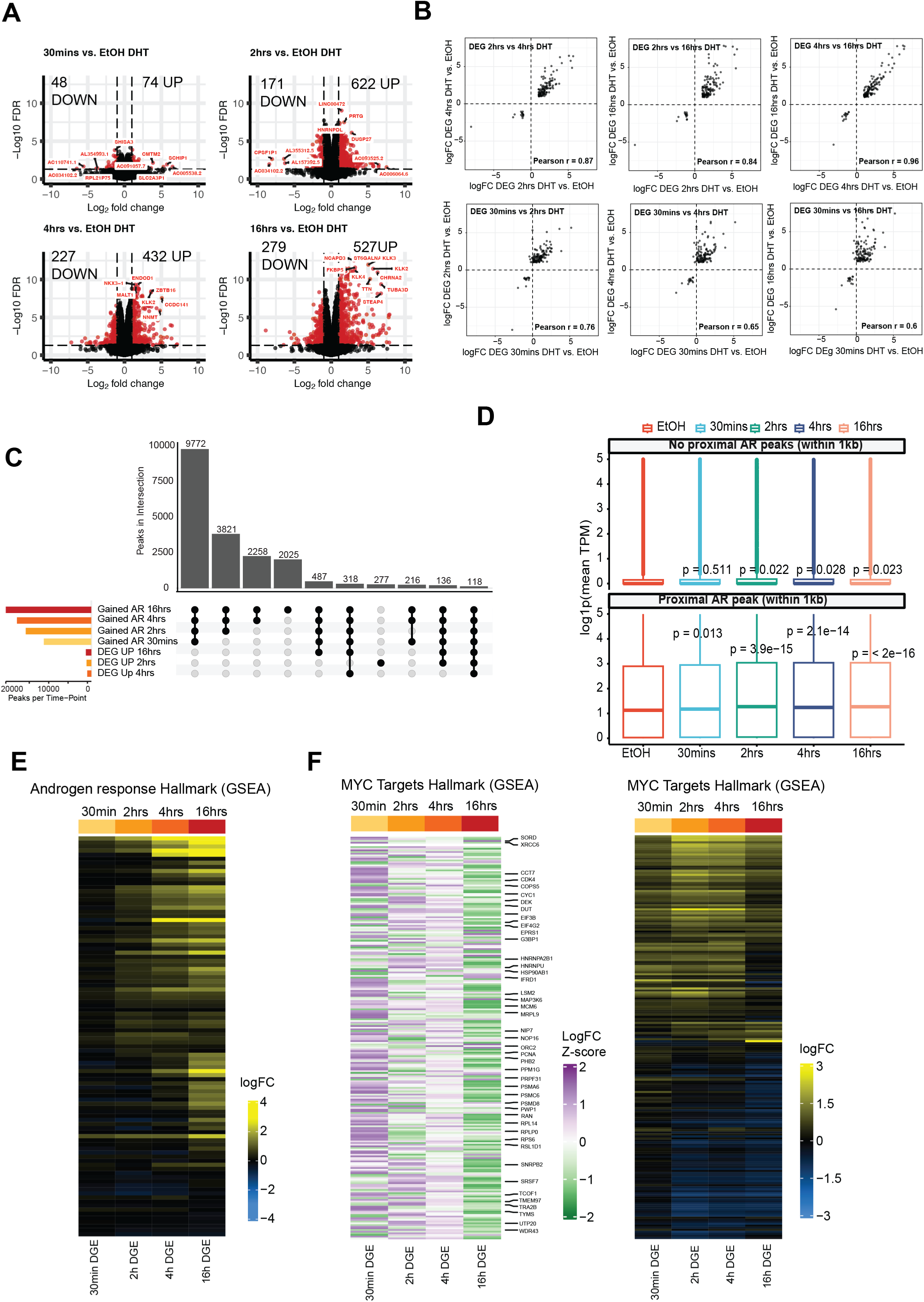
**(A)** Volcano plots (−log10(FDR) vs. log2 fold change) of all genes detected in mRNA differential gene expression analysis per DHT time-point compared to EtOH (red = significant; black = non-significant). **(B)** Correlation plots between log2 fold change (logFC) of differentially expressed genes (DEG) at each of the DHT treatment time-points. **(C)** Upset plot representing the overlap between differentially expressed genes at each DHT time-point compared to EtOH and gained AR binding sites at each DHT time-point compared to EtOH. **(D)** Box plot of gene expression (log TPM) for genes with (within 1Kb) and without a proximal promoter AR binding site (*P* value, Fisher’s exact test). **(E)** Heatmap showing change in expression (logFC vs. EtOH) of genes belonging to the Androgen Response (GSEA) Hallmark. **(F)** Heatmap showing change in expression (Z-scores of logFC vs. EtOH and logFC vs. EtOH) of genes belonging to the MYC Targets V1/V2 (GSEA) Hallmark.

**Supplementary Figure 3:**
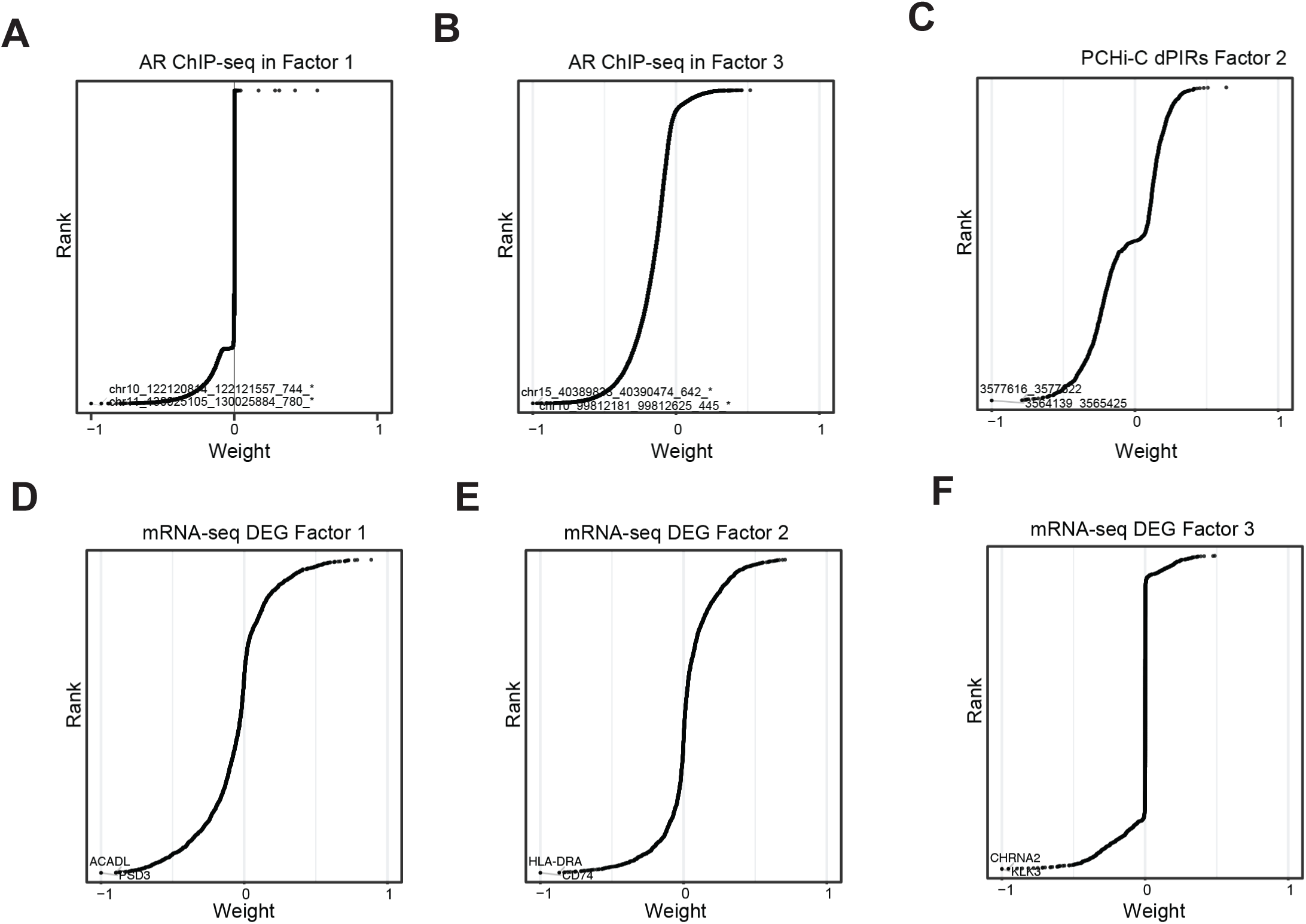
Overview and validation of the trained MOFA+ MEFISTO model. Inspection of feature weights shown by distribution plots ranked by loading weight assigned to each data point per latent factor, with the top 2 weighted datapoints labelled; **(A)** for AR ChIP-seq regions in Factor 1; **(B)** for AR ChIP-seq regions in Factor 3; **(C)** for PCHi-C dPIRs in Factor 2; **(D)** for mRNA-seq DEGs in Factor 1; **(E)** for mRNA-seq DEGs in Factor 2 and **(F)** for mRNA-seq DEGs in Factor 3.

**Supplementary Figure 4:**
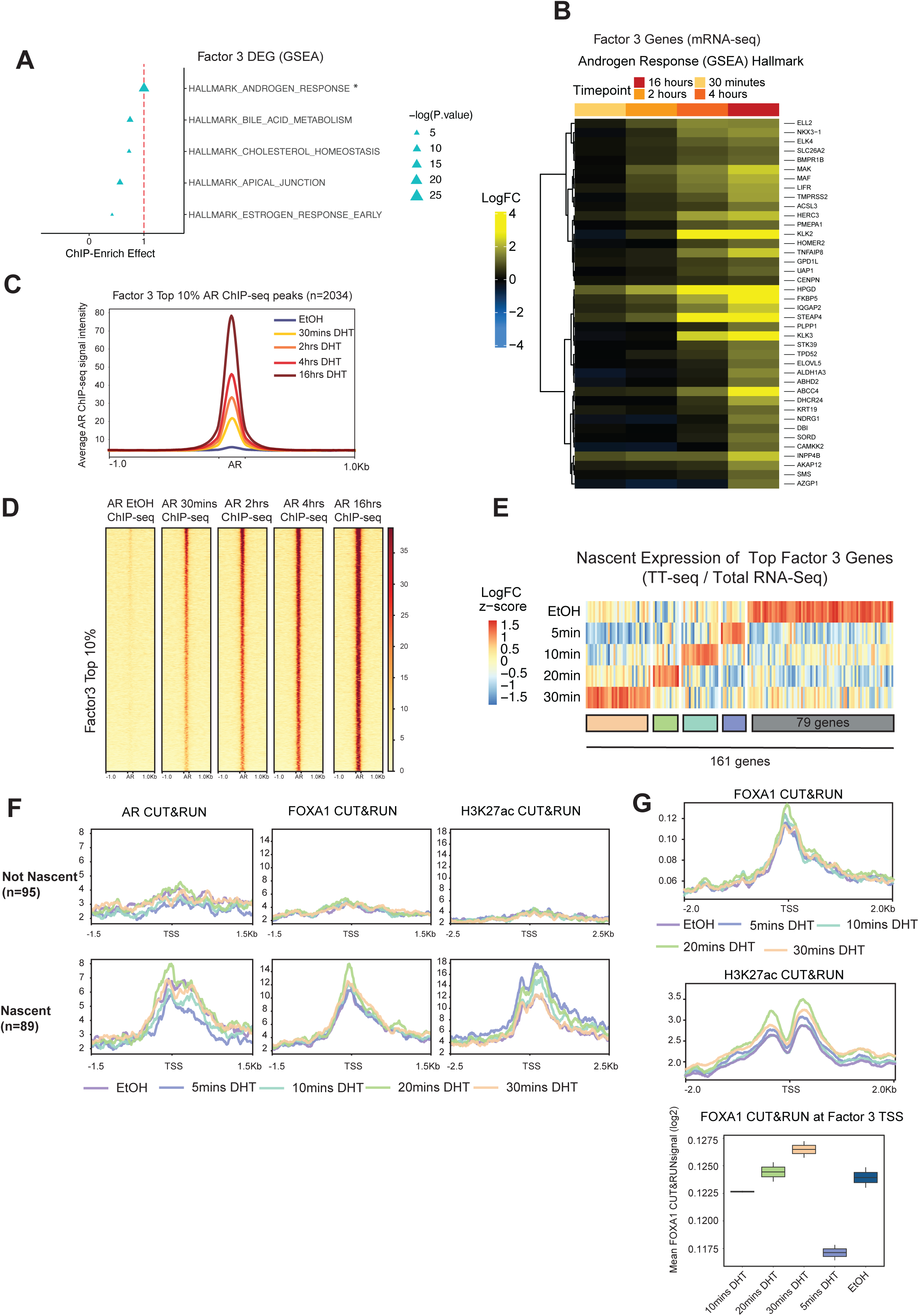
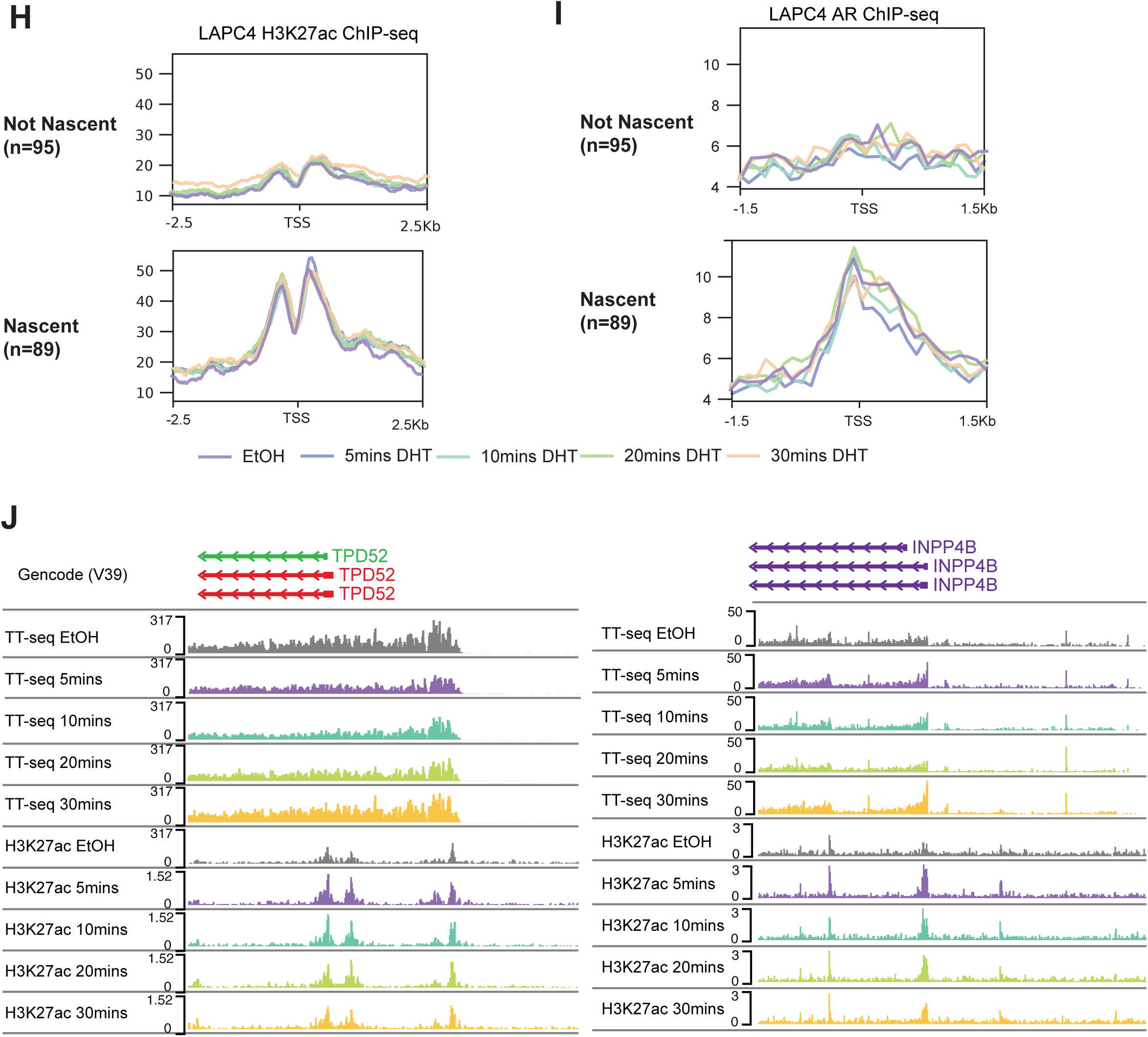
Characterization of the progressive transcription (Factor 3) subset. **(A)** Effect scores (ChIP-Enrich) for signature gene sets representing Factor 3 DEGs. Asterisk (*) indicates P value < 0.05 and effect score >1. **(B)** Heatmap showing change in expression (logFC vs. EtOH) of genes belonging to Factor 3 “Progressive Transcription” and Androgen Response (GSEA) Hallmark. **(C)** Average profiles of AR ChIP-seq signal intensity at top 10% AR ChIP-seq peaks in Factor 3 (“Progressive Transcription”) for each DHT time-point of and EtOH. **(D)** Heatmap of AR ChIP-Seq (n=3) signal intensity at top 10% AR ChIP-seq peaks in Factor 3 (“Progressive Transcription”) for each DHT time-point of and EtOH. **(E)** Heatmap showing nascent expression (Z-scores of logFC 4sU labelled TT-Seq vs. unlabelled Total RNA-Seq) across “early” DHT treatment time-course of genes in Factor 3 (“Progressive Transcription”). **(F)** Average CUT&RUN signal intensity for AR, FOXA1 and H3K27ac at nascently transcribed genes with maximal expression observed at each DHT treatment time-point. The top row represents the average signal intensity for non-nascently transcribed genes and the bottom row average signal intensity for nascently transcribed genes. **(G)** Average CUT&RUN signal intensity for FOXA1 and H3K27ac at all Factor 3 “Progressive Transcription” genes with maximal expression observed at each DHT treatment time-point. **(H)** Average LAPC4 ChIP-seq signal intensity for H3K27ac at nascently transcribed genes with maximal expression observed at each DHT treatment time-point. The top row represents the average signal intensity for non-nascently transcribed genes and the bottom row average signal intensity for nascently transcribed genes. **(I)** Average LAPC4 ChIP-seq signal intensity for AR at nascently transcribed genes with maximal expression observed at each DHT treatment time-point. The top row represents the average signal intensity for non-nascently transcribed genes and the bottom row average signal intensity for nascently transcribed genes. **(J)** Representative examples demonstrating the nascent expression of top weighted androgen response hallmark genes: *TPD52* (left panel) and *INPP4B* (right panel). H3K27ac CUT&RUN and TT-seq signal tracks (replicates merged) are shown.

**Supplementary Figure 5:**
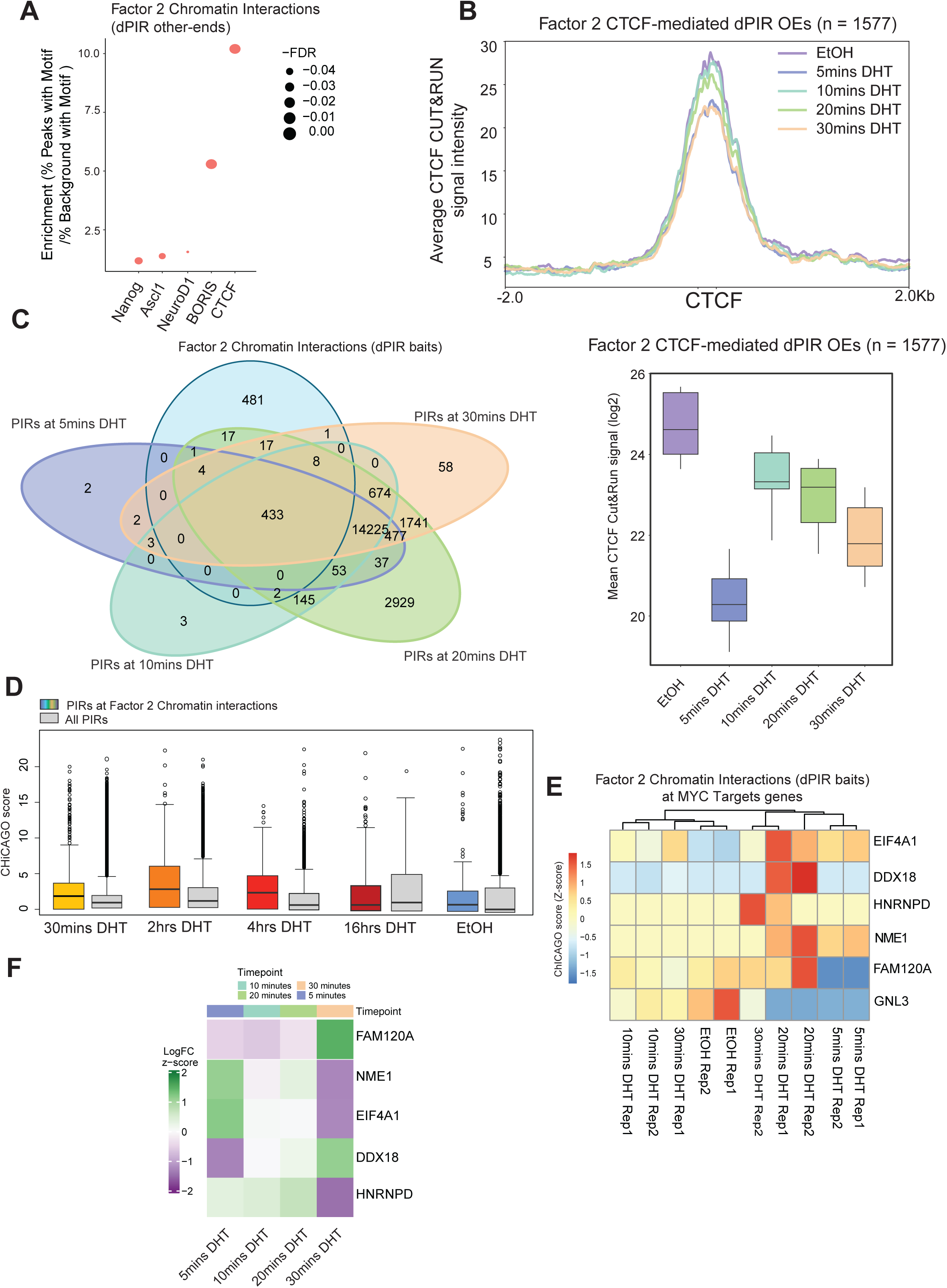
Characterization of the chromatin interactions (Factor 2) subset. **(A)** Transcription factor motifs enriched at top chromatin interaction (dPIRs other-ends) in Factor 2 plotted by FDR and percentage of peaks with motif (y-axis) compared to matched random background regions. **(B)** Average profiles of CTCF CUT&RUN signal intensity at CTCF-mediated dPIRs in Factor 2 (“Chromatin interactions”) for each time-point of the early DHT stimulation compared to EtOH control. Box plot quantification of the CTCF CUT&RUN average signal is provided below. **(C)** Venn diagram representing overlap between Factor 2 3D chromatin interactions (dPIR baits) and PIRs identified at each DHT time-point in the “early” DHT time-course. **(D)** Contact frequency (number of OEs per PIR) for Factor 2 (“Chromatin interactions”) dPIRs and all PIRs in the 16hrs DHT time-course. **(E)** Heatmap of CHiCAGO scores (Z-scores of weighted −log *P* values) for dPIRs belonging to top 10% Factor 2 regions at genes belonging to MYC Targets (V1/V2) (GSEA) Hallmark in the early DHT time-course. **(F)** Heatmap showing nascent expression (Z-scores of logFC 4sU labelled TT-Seq vs. unlabelled Total RNA-Seq) across “early” DHT treatment time-course of MYC Targets (V1/V2) Hallmark genes at Factor 2 3D chromatin interactions.

**Supplementary Figure 6:**
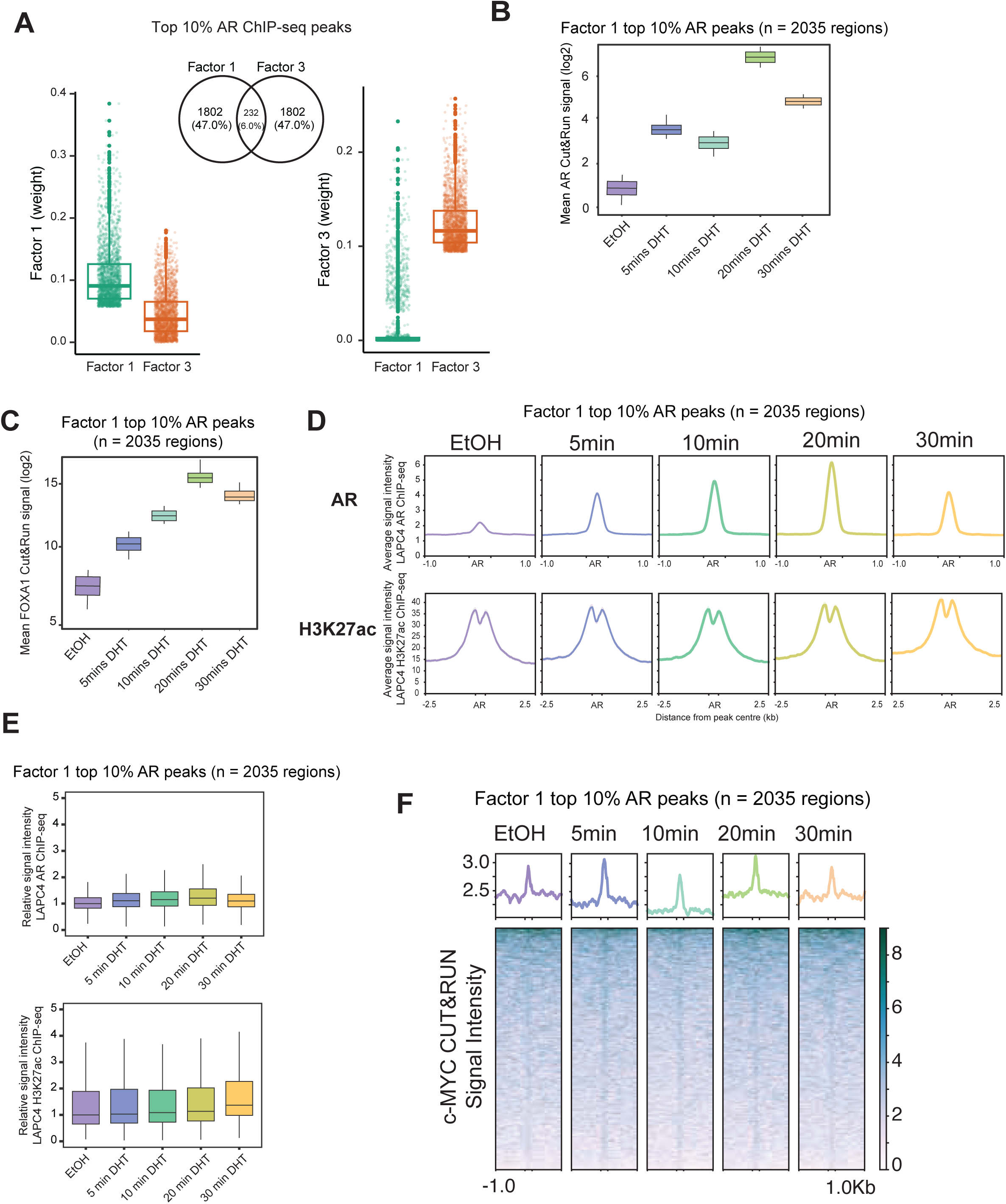
Characterization of the rapid AR binding (Factor 1) subset. **(A)** Box-and-whisker plot of absolute weighs assigned to the top 10% of AR peaks by Factor 1 (left panel) and Factor 3 (right panel). Venn Diagram (middle) shows the number of unique and common peaks in the top 10% of Factor 1 and Factor 3. **(B)** Box plot quantification of early DHT time-course AR CUT&RUN average signal profile at Factor 1 top 10% AR binding sites. **(C)** Box plot quantification of early DHT time-course FOXA1 CUT&RUN average signal profile at Factor 1 top 10% AR binding sites. **(D)** Average profiles of signal intensity for each time-point of the early DHT stimulation compared to EtOH control in LAPC4 cells; AR ChIP-seq (top panel) and H3K27ac ChIP-seq (bottom panel) at top 10% AR ChIP-seq peaks in Factor 1 (“Fast AR binding”). **(E)** Box plot quantification of early DHT time-course AR ChIP-seq (top panel) and H3K27ac ChIP-seq (bottom panel) average signal profile at Factor 1 top 10% AR binding sites. **(F)** Heatmap and average profiles of c-MYC CUT&RUN (n=3) signal intensity at top 10% AR ChIP-seq peaks in Factor 1 (“Fast AR binding”) for each time-point of DHT stimulation compared to EtOH control.

**Supplementary Figure 7:**
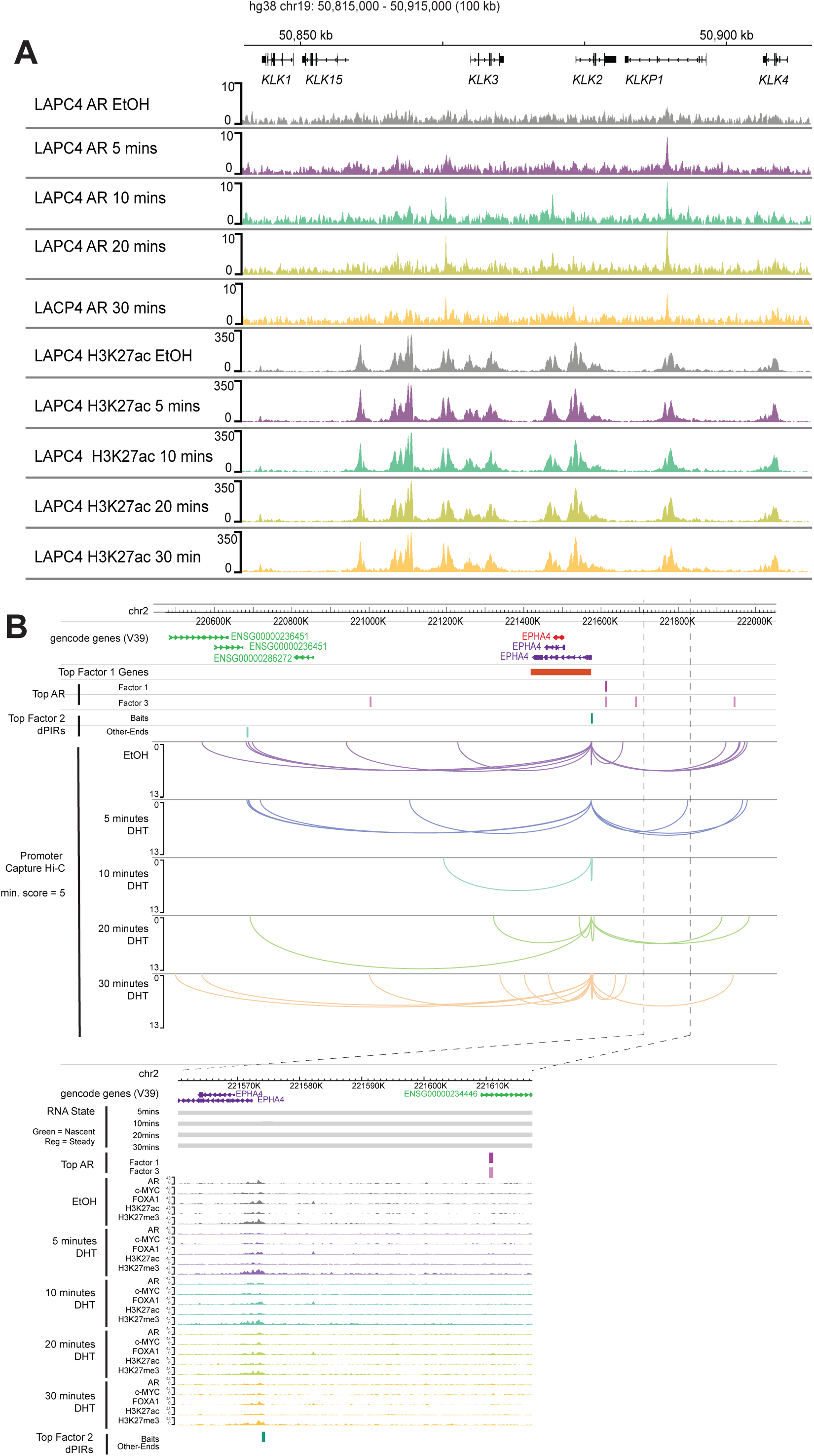
**(A)** Representative example of the *KLK* locus (*KLK3*, *KLK2* and *KLK4*) showing LAPC4 AR ChIP-Seq (n = 3) and H3K27ac ChIP-seq (n = 3) tracks. **(B)** Temporal dynamics of AR binding, transient 3D chromatin interactions and progressive transcription at the *EPHA4* locus. Representative example demonstrating the association “Fast” and “Progressive” AR binding sites (pink track) and “Progressive Transcription” gene expression (orange track) and “Transient Chromatin Interactions” (dPIR baits and OEs, teal track) as compared across the early DHT time-points. *EPHA4* locus shown. Nascent RNA state, AR, c-MYC, FOXA1, H3K27ac and H3K27me3 CUT&RUN and PCHi-C data (flat arc, zoomed-out 1.2Mb) tracks are shown.

**Supplementary Figure 8:**
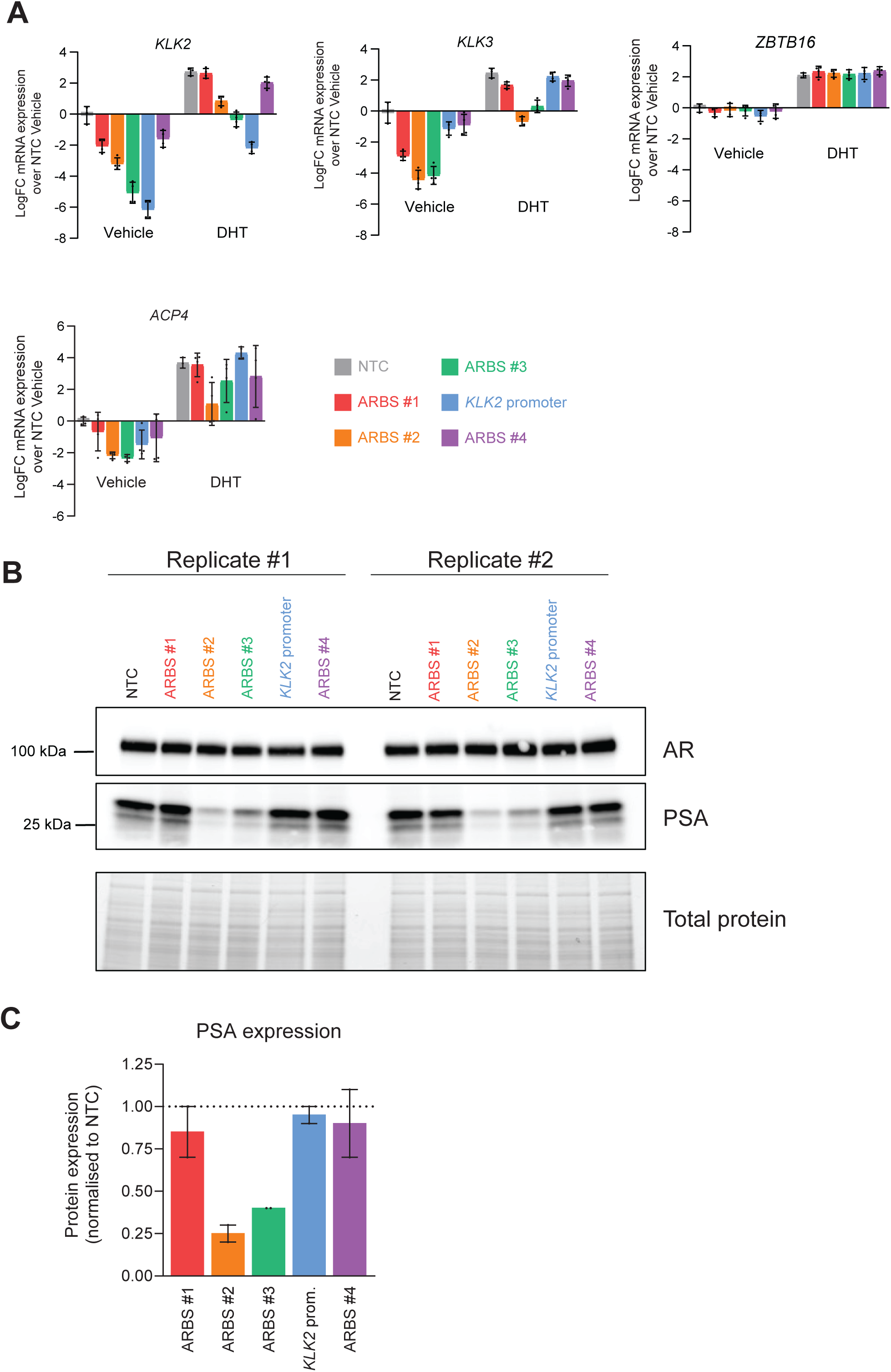
Androgen-mediated transcription is significantly reduced by inhibiting AR-bound enhancer regions. **(A)** Bar plots showing change in gene expression in CRISPRi compared with CRISPRi-empty controls (NTC) for genes located within the *KLK* locus. **(B)** Western blots for AR and PSA protein expression following 16hrs of DHT treatment in CRISPRi cells across two independent replicates. **(C)** Quantification of the protein expression in (**B**) using band densitometry.

## Notes

### Competing Interest Statement

The authors have declared no competing interest.

### Summary of Updates

We provide new validation experimental data sets and further analyses in an independent cell line models as well as functional studies.

